# Integrative Molecular Characterization of Sarcomatoid and Rhabdoid Renal Cell Carcinoma Reveals Determinants of Poor Prognosis and Response to Immune Checkpoint Inhibitors

**DOI:** 10.1101/2020.05.28.121806

**Authors:** Ziad Bakouny, David A. Braun, Sachet A. Shukla, Wenting Pan, Xin Gao, Yue Hou, Abdallah Flaifel, Stephen Tang, Alice Bosma-Moody, Meng Xiao He, Natalie Vokes, Jackson Nyman, Wanling Xie, Amin H. Nassar, Sarah Abou Alaiwi, Ronan Flippot, Gabrielle Bouchard, John A. Steinharter, Pier Vitale Nuzzo, Miriam Ficial, Miriam Sant’Angelo, Juliet Forman, Jacob E. Berchuck, Shaan Dudani, Kevin Bi, Jihye Park, Sabrina Camp, Maura Sticco-Ivins, Laure Hirsch, Megan Wind-Rotolo, Petra Ross-Macdonald, Maxine Sun, Gwo-Shu Mary Lee, Steven L. Chang, Xiao X. Wei, Bradley A. McGregor, Lauren C. Harshman, Giannicola Genovese, Leigh Ellis, Mark Pomerantz, Michelle S. Hirsch, Matthew L. Freedman, Michael B. Atkins, Catherine J. Wu, Thai H. Ho, W. Marston Linehan, David F. McDermott, Daniel Y.C. Heng, Srinivas R. Viswanathan, Sabina Signoretti, Eliezer M. Van Allen, Toni K. Choueiri

## Abstract

Sarcomatoid and rhabdoid (S/R) renal cell carcinoma (RCC) are highly aggressive tumors with limited molecular and clinical characterization. Emerging evidence suggests immune checkpoint inhibitors (ICI) are particularly effective for these tumors^1–3^, although the biological basis for this property is largely unknown. Here, we evaluate multiple clinical trial and real-world cohorts of S/R RCC to characterize their molecular features, clinical outcomes, and immunologic characteristics. We find that S/R RCC tumors harbor distinctive molecular features that may account for their aggressive behavior, including *BAP1* mutations, *CDKN2A* deletions, and increased expression of *MYC* transcriptional programs. We show that these tumors are highly responsive to ICI and that they exhibit an immune-inflamed phenotype characterized by immune activation, increased cytotoxic immune infiltration, upregulation of antigen presentation machinery genes, and PD-L1 expression. Our findings shed light on the molecular drivers of aggressivity and responsiveness to immune checkpoint inhibitors of S/R RCC tumors.

## Introduction

Sarcomatoid and rhabdoid (S/R) renal cell carcinoma (RCC) are among the most aggressive forms of kidney cancer^4, 5^. Sarcomatoid and rhabdoid features represent forms of dedifferentiation of RCC tumors and can occur in the same tumor or independently of each other^6^. These features can develop over any background RCC histology, including clear cell, papillary, and chromophobe RCC. These tumors account for 10-15% of RCC and most patients with S/R RCC present with metastatic disease^4, 7^. While classic RCC therapies such as VEGF and mTOR targeted therapies are largely ineffective for these tumors, multiple clinical studies suggest that immune checkpoint inhibitors (ICI) may have significant clinical activity in sarcomatoid and rhabdoid RCC^1–3, 8–11^. Prior studies have hinted that these tumors may harbor distinctive molecular features, although these studies were limited by small sample sizes, restricted molecular analyses, leading to discordant conclusions^2, 12–15^.

To define the molecular properties underlying the S/R clinical subtype and determine their relationship to potentially enhanced response to ICI, we perform an expanded clinical and molecular integrated characterization of S/R RCC in both clinical trial and real-world cohorts, assessing clinical outcomes on ICI, genomic and RNA sequencing (RNA-seq), immunohistochemical (IHC) staining for PD-L1, immunofluorescence (IF)-based assessment of immune infiltration, and transcriptomic evaluation of sarcomatoid cell lines (Fig. 1a).

**Figure 1:**
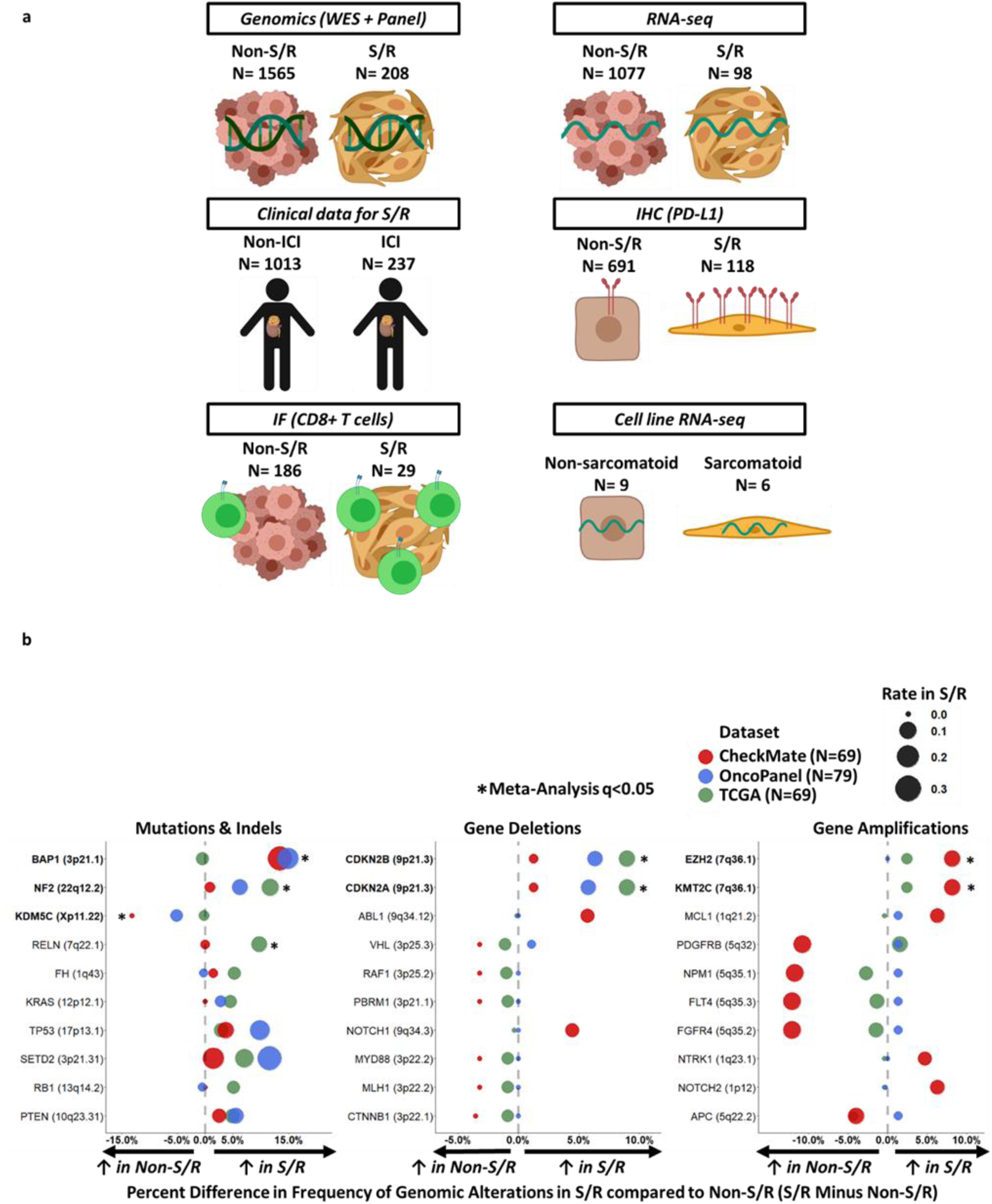
Genomic characterization of S/R RCC reveals distinctive genomic features. (a) Overview of the clinical, molecular, and cell line data. (b) Comparison of S/R vs. non-S/R RCC by mutations & indels, deep deletions, and high amplifications in the CheckMate, OncoPanel, and TCGA cohorts. *q<0.05 (Fisher’s method meta-analysis of Fisher’s exact tests); ICI: Immune Checkpoint Inhibitor; IF: Immunofluorescence; IHC: Immunohistochemistry; RNA-seq: RNA-sequencing; S/R: Sarcomatoid/Rhabdoid; TCGA: The Cancer Genome Atlas; WES: Whole Exome Sequencing

## Results

### S/R RCC Tumors Harbor Distinctive Genomic Features

We first evaluated the genomic landscape of S/R RCC (total N= 208) in three distinct cohorts (two whole exome sequencing [WES] and 1 gene panel sequencing cohort [OncoPanel]) and compared it to that of non-S/R RCC (total N= 1565; Table S1). This DNA-sequencing cohort included one clinical trial WES cohort (CheckMate cohort), a retrospective analysis of an institutional panel-based sequencing cohort (OncoPanel cohort), and a retrospective pathologic review and analysis of a publicly available cohort (TCGA cohort). The most commonly altered genes in S/R RCC (Fig. S1) were generally similar to those previously reported for RCC^16^. We subsequently compared the genomic features of S/R RCC tumors to background histology-matched non-S/R RCC tumors across the three cohorts. Tumor mutational burden (TMB), total indel load, and frameshift indel load were overall similar between S/R RCC and non-S/R RCC tumors (Fig. S2a-c). While the frameshift indel load was significantly increased (p= 0.024) in S/R vs. non-S/R RCC in the OncoPanel cohort, the absolute difference was small and was not corroborated in the two WES cohorts (CheckMate and TCGA; Fig. S2c).

Next, gene-specific alteration rates were compared between S/R and non-S/R RCC in each of the three cohorts independently and in combination (Methods). *BAP1* and *NF2* somatic alterations were significantly and consistently enriched in S/R compared to non-S/R RCC, whereas *KDM5C* somatic alterations were significantly less frequent in S/R compared to non-S/R RCC (Fisher’s exact q<0.05; Fig. 1b and Table S2). Furthermore, *CDKN2A* and *CDKN2B* deep deletions as well as *EZH2* and *KMT2C* high amplifications were significantly enriched in S/R compared to non-S/R (Fisher’s exact q<0.05 and consistent across at least two of the three included datasets; Fig. 1b and Table S2). Other genes that were significantly amplified (low or high amplification) included *MYC* and *CCNE1,* whereas those that were significantly deleted (shallow or deep deletion) included *RB1* and *NF2* (Fisher’s exact q<0.05). Although recent reports have suggested that genes in the 9p24.1 locus (including *CD274*, *JAK2*, and *PCD1LG2* genes) were more frequently amplified in RCC tumors with sarcomatoid features^2, 17^, we did not observe focal amplifications to be enriched at this locus (Table S2). Moreover, differences between S/R and non-S/R RCC were generally consistent regardless of background histology (clear cell or non-clear cell; Table S2).

Since the analyses in this study are based on single region sampling of S/R RCC tumors and since such sampling has been shown to affect the detection rate of mutations in RCC tumors^18^, we next compared the intra-tumoral heterogeneity (ITH) index between S/R and non-S/R RCC tumors (Methods). We found that the ITH index was not significantly different between these two groups of tumors in the CheckMate cohort. Furthermore, this was corroborated in a re-analysis of the TRACERx Renal study, whereby the ITH index did not differ between S and non-S RCC tumors (Fig. S3a). Moreover, among 71 S/R RCC tumors in the OncoPanel cohort (of a total of 79 S/R RCC tumors) for which the portion of the tumor that was sequenced was assessable, 44 tumors had the S/R (mesenchymal) regions sequenced and 27 had the non-S/R (epithelioid) regions of the tumor sequenced. These two subsets of tumors were compared and no significant overall mutation/indel load (Fig S3b) or gene-level mutational (Table S3) differences were found, other than a marginal but statistically significant (p= 0.042) increase in the number of frameshift indels in mesenchymal regions. In addition, panel sequencing mutation data from 23 sarcomatoid tumors that had been laser micro-dissected (into sarcomatoid and epithelioid components) and sequenced separately from the study by Malouf et al.^19^ was re-analyzed. In accordance, with the above findings no significant overall mutation/indel load (Fig S3c) or gene-level mutational (Table S3) differences were found. However, it should be noted that alteration frequency for certain genes differed between mesenchymal and epithelioid portions of S/R RCC tumors (Table S3). While certain mutations may be enriched in these tumors (in particular TP53 mutations, as has been previously suggested^14^), none rose to the level of statistical significance in our cohort. Overall, our results suggest that the mutational differences between S/R and non-S/R RCC tumors are more pronounced than intra-tumoral mutational differences between mesenchymal and epithelioid portions of a given S/R RCC tumor. S/R RCC tumors have a distinctive genomic profile characterized by an enrichment for genomic alterations previously associated with poor prognosis in RCC (such as *BAP1* and *CDKN2A*) and genomic alterations that may represent therapeutic targets in S/R RCC (*CDKN2A* and *CDKN2B* deletions, *EZH2* amplifications, and *NF2* mutations).

### Transcriptomic Programs of S/R RCC Underpin their Poor Prognosis

We next assessed transcriptomic programs in S/R RCC and their relationship to the known poor prognosis of this subtype. We compared RNA-seq data between S/R (total N= 98) and non-S/R RCC (total N= 1076) in the TCGA (publicly available) and CheckMate cohorts independently (Methods; Table S4) using Gene Set Enrichment Analysis (GSEA)^20^. Twelve gene sets were upregulated (GSEA q<0.25) in S/R compared to non-S/R RCC in the two cohorts independently, including cell cycle programs, genes regulated by *MYC*, and apoptosis programs (Fig. 2a; Table S5). Specific upregulated gene sets may account for their morphological features including their mesenchymal appearance^6^ (upregulation of epithelial-mesenchymal-transition [EMT]) and frequent co-occurrence of necrosis (endoplasmic reticulum [ER] stress and apoptosis-caspase pathway)^4, 7^, and rapid progression (E2F targets, G2/M checkpoint, mitotic spindle assembly). Moreover, high *MYC* targets version 1 (v1) expression as quantified by single sample GSEA (ssGSEA) scores^21^ significantly correlated with worse clinical outcomes in both the subset of patients with S/R in the anti-PD-1 (nivolumab) arm of the CheckMate cohort as well as the subgroup of stage IV S/R RCC patients in TCGA independently (Fig. 2b; Fig. S4; Table S6). Of note, the majority of founder gene sets of both the *MYC* v1 and v2 “Hallmark” gene were enriched in S/R RCC (Fig. S5a), further corroborating the fact that *MYC*-regulated transcriptional programs are enriched in S/R RCC. Moreover, the correlation with outcomes within S/R RCC of the *MYC* v1 score was consistent when the *MYC*-regulated transcriptional program was measured using the separate but related *MYC* v2 “Hallmark” gene set (Fig. S5b-c). Patients with non-S/R RCC and *MYC* v1 scores similar to those of S/R RCC (above the median of the S/R RCC group for *MYC* v1) had significantly worse outcomes in both the TCGA and CheckMate PD-1 cohorts (Fig. 2c; Fig. S4; Table S6). These results indicate that a *MYC*-driven transcriptional program is driving the aggressive phenotype of S/R RCC tumors (also shared with a subset of non-S/R RCC)^5^.

**Figure 2:**
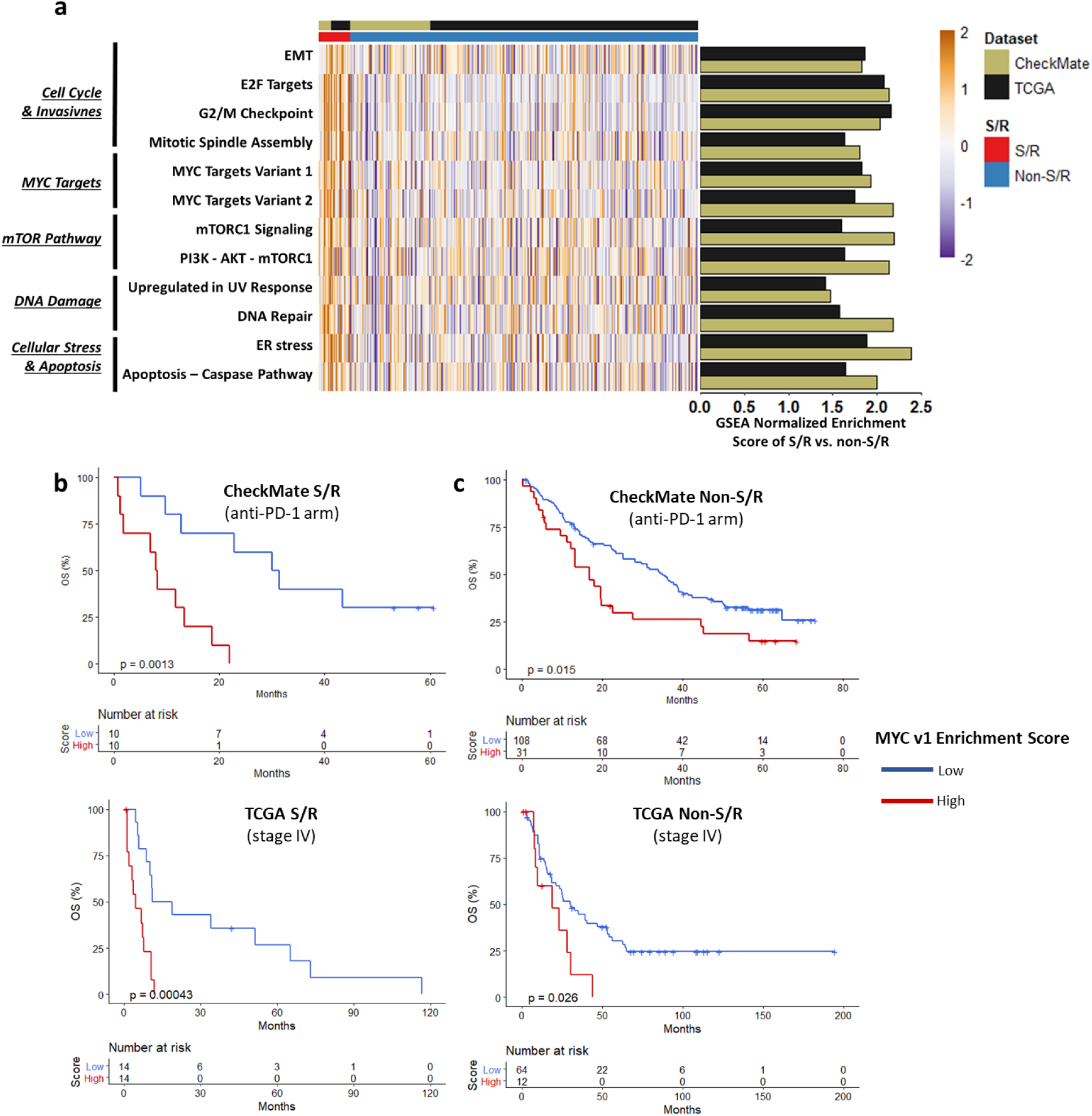
Transcriptional profiling of S/R RCC reveals the molecular correlates of its poor prognosis and identifies subsets of non-S/R tumors associated with a poor prognosis. (a) Heatmap and bar plots of the ssGSEA scores and GSEA normalized enrichment scores for the non-immune “Hallmark” gene sets that were found to be significantly enriched (q<0.25) in S/R compared to non-S/R RCC in both the TCGA and CheckMate cohorts independently. (b) Kaplan-Meier curves for OS by *MYC* v1 score within the S/R group of the CheckMate (anti-PD-1 arm) and TCGA (stage IV) cohorts; *MYC* v1 score dichotomized at the median. (c) Kaplan-Meier curves for OS by *MYC* v1 score within the non-S/R group of the CheckMate (anti-PD-1 arm) and TCGA (stage IV) cohorts; *MYC* v1 score dichotomized at the median of the S/R group. EMT: Epithelial Mesenchymal Transition; *MYC* v1: *MYC* Targets Version 1; S/R: Sarcomatoid/Rhabdoid; TCGA: The Cancer Genome Atlas

Extending from the Hallmark GSEA analysis, 243 genes had significantly increased expression in S/R compared to non-S/R RCC independently across the two cohorts, including multiple cell cycle and proliferation (*CCNB1, CDC45, CDC6, CDCA3, CDCA7, CDCA8, CDK6,* and *MKI67*), immune *(HIVEP3, IFI16, IFI35, IL15RA, and LAG3),* and metastasis-implicated^22^ (*ACTB, ANLN, ARPC1B, ARPC5,* and *ARPC5L, CD44)* genes as well as chemokine (*CXCL9*) and antigen presenting machinery (*TAP1, TAP2, CALR, PSMA5, PSMB10, PSMB4, PSMC2, PSME2*) genes that may be driving the immune infiltration in these tumors (Table S7). Since the overexpression of antigen presentation machinery genes has been found to correlate with increased cytotoxic immune infiltration and ICI responsiveness^23^, we further explored the antigen presentation machinery genes using four dedicated REACTOME^24^ and KEGG^25^ gene sets and found all four to be significantly increased in both the CheckMate and TCGA cohorts independently (Table S5). In addition, 83 genes had significantly decreased expression including cell junction-implicated (*TJP1* and *DSC2*) and cell differentiation genes (*MUC4*; Table S7).

### S/R RCC Tumors Display Marked Sensitivity to Immune Checkpoint Inhibitors and an Immune-Inflamed Phenotype

With the unique molecular background of S/R RCC defined, we then sought to establish whether S/R RCC patients treated by immune checkpoint inhibitors (ICI) had improved clinical outcomes, as suggested by early studies, and whether particular molecular features established the basis for such clinical phenotypes. Patients with S/R RCC had improved outcomes on ICI compared to non-ICI agents across 3 cohorts (total N ICI arms = 237; total N non-ICI arms = 1013; Table S8): a local Harvard cohort, the multicenter International Metastatic RCC Database Consortium (IMDC) cohort, and a pooled analysis of the S/R subgroup of 2 clinical trials (CheckMate 010^26^ and CheckMate 025^27^) evaluating an anti-PD-1 agent (nivolumab) for metastatic RCC. Patients with S/R RCC had significantly improved outcomes on ICI compared to non-ICI across cohorts and clinical outcomes including overall survival (OS), progression free survival (PFS), time to treatment failure (TTF), and objective response rate (ORR; Fig. 3a-c).

**Figure 3:**
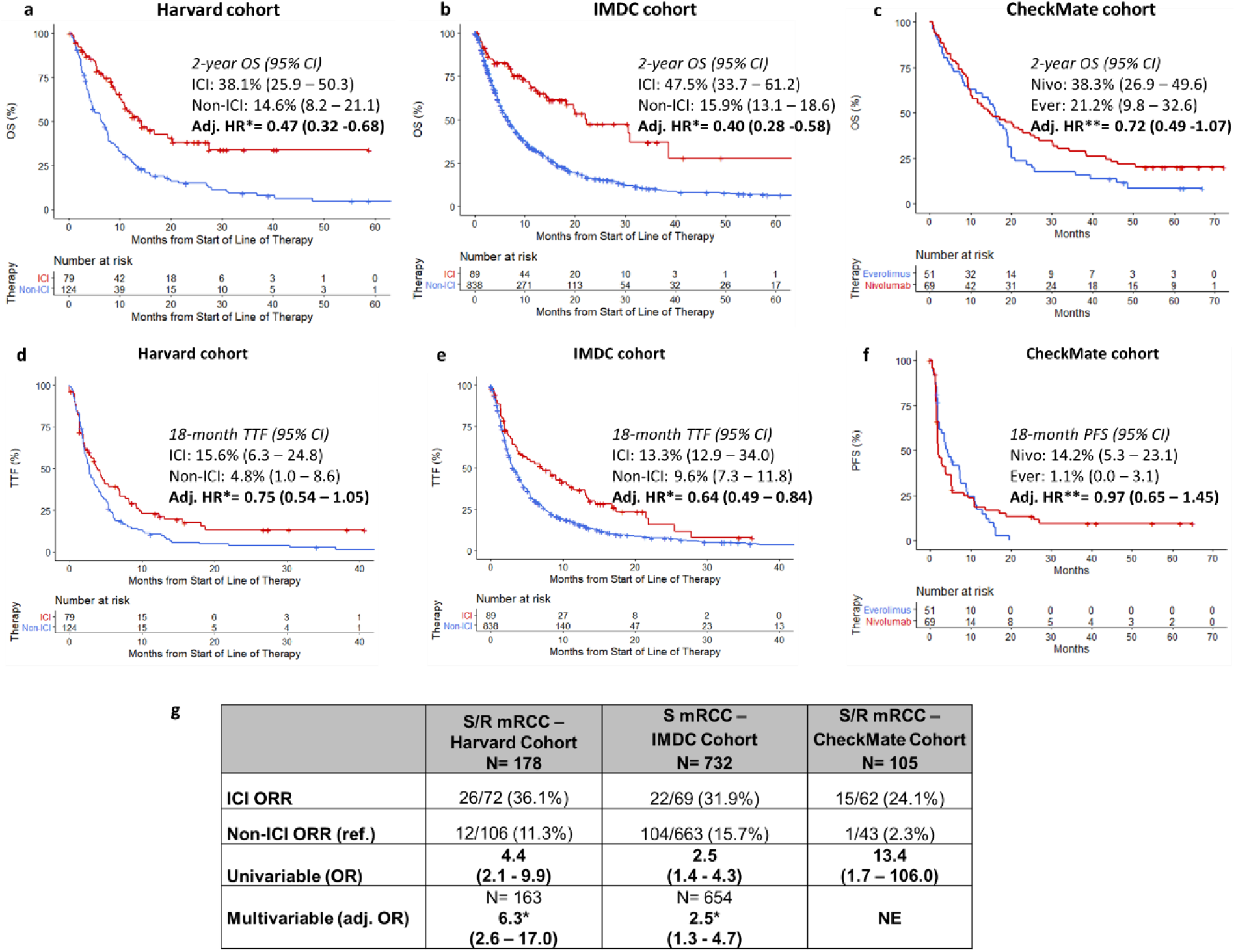
Improved clinical outcomes of S/R RCC tumors on immune checkpoint inhibitors across clinical trial and real-word cohorts. OS on ICI compared to non-ICI in the (a) Harvard, (b) IMDC, and (c) CheckMate S/R RCC cohorts. TTF on ICI compared to non-ICI in the (d) Harvard and (e) IMDC S/R RCC cohorts, and (f) PFS in the CheckMate S/R RCC cohort. (g) Summary table of overall response rate (among evaluable patients) on ICI compared to non-ICI in patients with S/R RCC across the Harvard, IMDC, and CheckMate cohorts. 95% CI: 95% Confidence Interval; Adj. Adjusted; Ever: Everolimus; HR: Hazard Ratio; ICI: Immune Checkpoint Inhibitor; IMDC: International Metastatic Renal Cell Carcinoma Database Consortium; Nivo: Nivolumab; NE: Not Evaluable; OS: Overall Survival; S/R: Sarcomatoid/Rhabdoid. * Adjusted for IMDC (International Metastatic Renal Cell Carcinoma Database Consortium) risk groups, line of therapy, and background histology. ** Adjusted for MSKCC (Memorial Sloan Kettering Cancer Center) risk groups

Given the significant sensitivity of S/R RCC to ICI as reflected by improved responses and survival outcomes, we examined molecular features that may drive this phenotype. First, GSEA on the immune “Hallmark” gene sets of the RNA-seq data of the TCGA and CheckMate cohorts showed that all 8 “Hallmark” immune gene sets were enriched (GSEA q<0.25) in S/R compared to non-S/R RCC in the two cohorts independently (Fig. 4a; Table S4), including gene sets previously implicated in response to ICI (e.g. interferon gamma response)^28, 29^. We then inferred immune cell fractions using the CIBERSORTx deconvolution algorithm (total N of S/R= 97 and Total N of non-S/R= 1028) and previously described gene signatures for Th1, Th2, and Th17 cells^30^ on the RNA-seq data from the CheckMate and TCGA cohorts. CD8+ T cell infiltration, CD8+/CD4+ T cell ratio, activated/resting NK cell ratio, M1 macrophages, M1/M2 macrophage ratio, as well as the Th1 score were all significantly increased (Mann-Whitney q<0.05) in S/R RCC in both cohorts independently (Fig. 4b, Fig S6a; Table S9). Moreover, the transcriptomic and immune microenvironment features of S/R RCC were consistent across S/R RCC subtypes (rhabdoid, sarcomatoid, or sarcomatoid and rhabdoid; Fig. S7-9).

**Figure 4:**
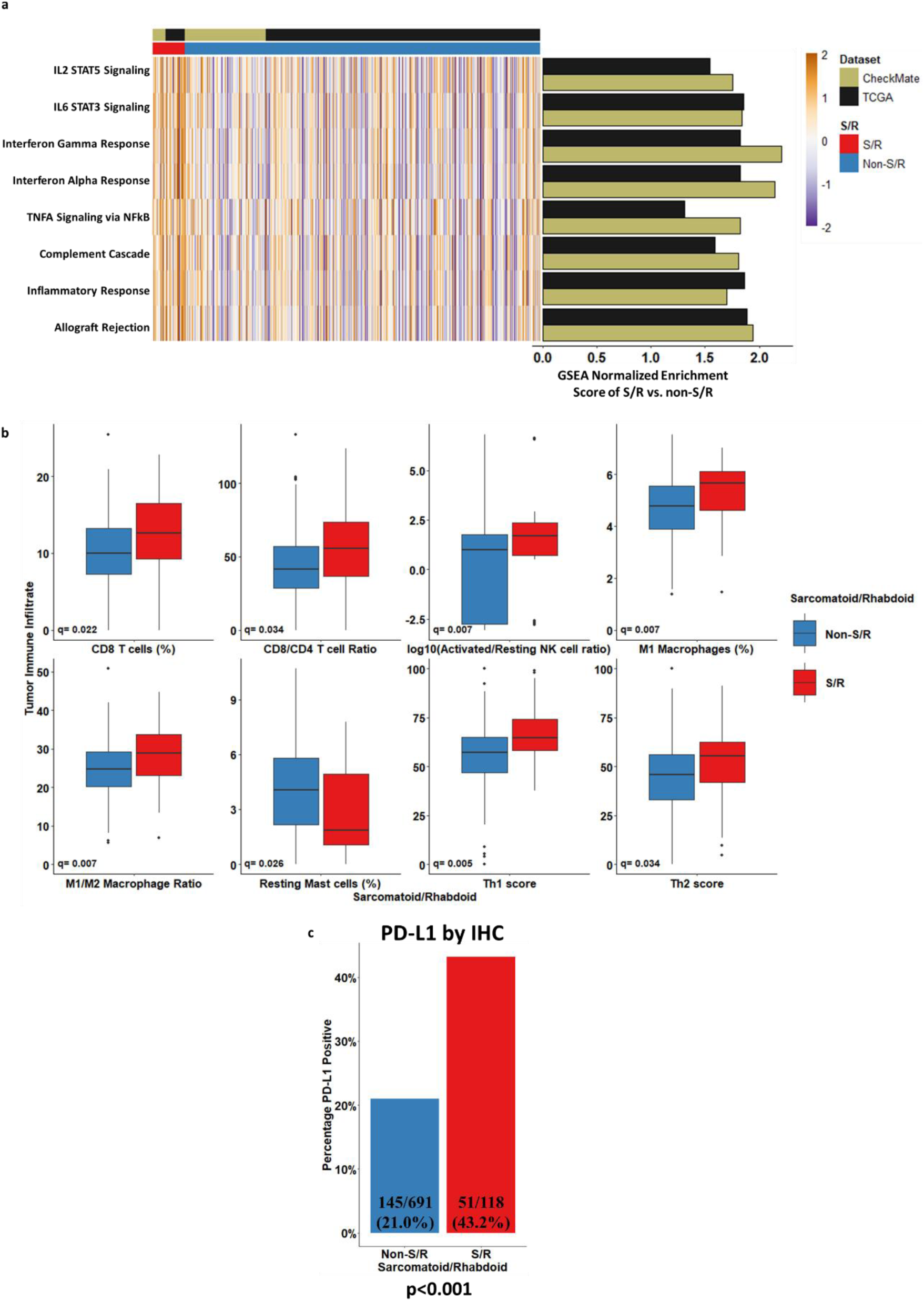
(a) Heatmap and bar plots of the ssGSEA scores and GSEA normalized enrichment scores for the immune “Hallmark” gene sets that were found to be significantly enriched (q<0.25) in S/R compared to non-S/R RCC in the TCGA and CheckMate cohorts independently. (b) Boxplots of the comparison of CIBERSORTx and Th immune cell populations between S/R and non-S/R RCC, with Mann-Whitney U test comparisons corrected for multiple comparison testing (q value reported). Only variables which were significant (q<0.05) in both the CheckMate and TCGA cohorts independently are shown. The CheckMate results are displayed in this figure. (c) Bar plot of the comparison of the proportions of tumors that were PD-L1 positive (≥1% on tumor cells) in S/R compared to non-S/R RCC. Fisher’s exact test p-value reported. TCGA: The Cancer Genome Atlas.

The immune-inflamed phenotype of S/R RCC tumors was further corroborated by an immunohistochemistry (IHC; N of S/R= 118 and N of non-S/R= 691) assay showing significantly increased PD-L1 (cut-off of ≥1%) expression on tumor cells in S/R compared to non-S/R tumors (43.2% vs. 21.0%; Fisher’s exact p<0.001; Fig 4c and Table S10) in the CheckMate cohort. To evaluate whether the elevated PD-L1 expression in S/R RCC is driven by PD-L1 gene amplification, as previously reported^2, 17^, we compared IHC-based PD-L1 expression by *CD274* (or PD-L1) gene copy number status (N= 63 patients in the S/R CheckMate cohort). We found that S/R tumors had increased PD-L1 expression (relatively to non-S/R RCC) independent of *CD274* copy number status (any deletion, amplification, or neither; all deletions were one-copy deletions); although the three S/R patients with *CD274* gene amplification (1 patient with high amplification and 2 with low amplifications) all expressed PD-L1 by IHC above the cut-off of ≥1%. Moreover, *CD274* copy number status did not correlate with clinical outcomes in patients treated with a PD-1 inhibitor (Fig. S10a-c). The immune-inflamed phenotype of S/R RCC tumors was also evaluated by IF staining for CD8+ T cells in a subset of the CheckMate cohort (N of S/R= 29 and N of non-S/R= 186; Fig S6b-c and Table S10). CD8+ T cell infiltration at the tumor invasive margin, which had been reported to be associated with response to ICI-based therapies^31^, tended to be increased in these tumors (although the difference was not statistically significant, Mann-Whitney p= 0.14). Since *BAP1* mutations are enriched in S/R RCC tumors in this study and have been previously associated with immune infiltration and inflammation^32^, we evaluated whether the immune findings reported in this study are driven by *BAP1* mutations. In a sensitivity analysis excluding all *BAP1* mutants (from the S/R and non-S/R RCC) groups, the immune findings reported in this study were found to be largely consistent with the results of the primary analysis, suggesting that the immune findings of the current study in S/R RCC tumors are not driven by *BAP1* mutations (Fig. S11). Taken together, S/R RCC tumors are highly responsive to ICI-based therapies and an immune-inflamed microenvironment in S/R RCC may be driving these responses in a *BAP1*-independent manner, leading to improved survival on ICI.

### Sarcomatoid Cell Lines Recapitulate the Biology of S/R RCC tumors

To evaluate which transcriptomic programs enriched in S/R RCC tumors were attributable to sarcomatoid cancer cells rather than the microenvironment, we compared baseline RNA-seq data from 6 distinct sarcomatoid kidney cancer cell lines and 9 distinct non-sarcomatoid kidney cancer cell lines (Fig S12a-b; Table S11). The transcriptional profile observed from the bulk profiling of tumors was partially recapitulated in the cell lines, with EMT and apoptosis-caspase pathway genes significantly enriched in sarcomatoid cell lines compared with non-sarcomatoid cell lines (Fig S12b). Given the shared transcriptional programs between sarcomatoid tumors and cell lines, we then sought to nominate candidate pathways that might reflect selective dependencies of sarcomatoid tumor cells. For this exploratory analysis, we interrogated publicly available data from 20 kidney cancer cell lines with both baseline RNA-seq and cell line drug response data. Among this group of 20 kidney cancer cell lines screened with 437 compounds of diverse mechanisms of action, we found EMT and apoptosis-caspase pathway ssGSEA scores most strongly correlated with sensitivity to cyclin dependent kinase inhibitors (CDKi; Fig. S12c; Table S11) and compared favorably to other classic therapeutic targets in RCC such as VEGF and mTOR inhibitors, consistent with the poor response of S/R RCC tumors to these agents^5, 33^. In an attempt to corroborate these findings we focused on two CDKi agents, SNS-032 and alvocidib, that displayed a strong correlation of their sensitivity profiles with the EMT and apoptosis-caspase signature scores in CTRP (Fig. S12; Fig. S13a-b; Table S11). In an independent in silico analysis of the recently published PRISM cell line drug screen dataset^34^, a similar relationship between sensitivity to CDKi and the EMT and apoptosis signatures was found for alvocidib and other CDKi (Fig. S13a; Table S10; SNS-032 was not tested in the PRISM dataset). SNS-032, alvocidib, and a VEGF inhibitor control agent (axitinib) were also separately evaluated in two sarcomatoid RCC cell lines (UOK127 and RCJ41-T2; not included in the CTRP or PRISM screens) and three non-sarcomatoid RCC cell lines (Caki-2, KMRC-20, and KMRC-2; included in the CTRP or PRISM screens). Although the relative sensitivities for the non-sarcomatoid cell lines determined in CTRP/PRISM globally mirrored relative sensitivities upon validation, we did not observe marked differential sensitivity between sarcomatoid and non-sarcomatoid cell lines for any of the 3 agents tested (Fig S14).

## Discussion

The current study represents a large integrative molecular and clinical characterization of S/R RCC, including clinical outcomes on ICI therapies and non-ICI controls from both clinical trial and retrospective cohorts, DNA and RNA-sequencing data, IHC and IF-based assessment of the immune microenvironment, and the molecular profiling of cell line models of the disease. We show that S/R RCC tumors are highly responsive to ICIs, harbor distinctive genomic alterations, a characteristic transcriptional program characterized by the enrichment of *MYC*-regulated genes that correlates with poor outcomes, and a heavily inflamed microenvironment enriched in features that have been associated with ICI responses.

Our genomic findings corroborate those of prior studies that reported significant enrichment of Hippo pathway (which includes the *NF2* gene) mutations^19^ in S vs non-S RCC tumors and *BAP1* mutations in S and R RCC tumors^12, 15, 35^. While *CDKN2A* alterations have been reported in S RCC tumors^13, 19^, these alterations are also present in non-S/R RCC tumors^36^. However, the current study established *CDKN2A/B* deep deletions as specifically enriched in S/R compared to non-S/R RCC tumors as well as depletion in *KDM5C* mutations and enrichment in *EZH2* amplifications in S/R RCC tumors. Moreover, S/R RCC tumors were not found to consistently harbor a significantly increased rate of mutations, indels, or frameshift indels compared with non-S/R RCC tumors.

S/R RCC tumors are rapidly proliferating tumors that are associated with poor prognosis and rapid clinical progression^37, 38^. While prior studies had identified multiple clinical and pathological factors that are associated with prognosis in patients with S/R RCC tumors^39, 40^, the molecular drivers of aggressivity of S/R RCC tumors had largely been unexplored. Here, we show that multiple molecular pathways implicated in cell cycle regulation and invasiveness as well as *MYC*-regulated genes are enriched in S/R RCC tumors and that the enrichment in *MYC*-regulated genes correlates with poor prognosis. These results suggest that *MYC*-regulated transcriptional programs are key factors driving the aggressivity and poor prognosis associated with S/R RCC tumors.

While prior studies have largely reported on tumors with sarcomatoid features, the different cohorts of this study highlight that rhabdoid features frequently co-occur with sarcomatoid features (10-20% of S/R RCC tumors). In addition, tumors harboring rhabdoid features alone are also relatively frequent (5-25% of S/R RCC tumors). In this study, the molecular features of S, R, and S+R (harboring both features concurrently) tumors were not found to be significantly different (Figure S1 and Figures S7-S9). However, detecting smaller effect sizes in these comparisons was limited by the relatively small sample sizes of the R and S+R groups.

The preliminary clinical outcomes of the subgroups of patients with S RCC from four large randomized clinical trials of the first line treatment of metastatic RCC ^8–11^ reported ORRs ranging between 46.8% and 58.8% for patients with S RCC treated with first line ICI combinations, with a significant clinical benefit compared to the non-ICI control arms (sunitinib in all four trials). These results for ICI arms are numerically superior to those reported in the current study (ORR range 24.1-36.1% in ICI arms). Multiple potential factors could account for the increased effectiveness observed in these preliminary reports of subgroup analyses of phase III randomized controlled trials, compared to the findings in the three cohorts included in the current study. Indeed, the ICI arms in these studies were combination therapies (either PD-1 inhibitor + CTLA-4 inhibitor or PD-(L)1 + VEGF inhibitor) and all patients were being treated in the first line setting (and therefore not previously refractory to other therapies). In the current study, patients with S/R RCC derived significant clinical benefit from ICI regimens while having been treated by various different ICI regimens (entirely ICI monotherapy in the CheckMate cohort and with a large proportion of ICI monotherapy in the IMDC and Harvard cohorts; Table S7) and across different lines of therapy in each of the three cohorts (with a substantial proportion in the second line and beyond). Our findings, derived from three independent cohorts, suggest that S/R RCC tumors derive benefit from ICI regimens even outside of the setting evaluated in the subgroup analyses of the above-mentioned phase III trials (first line ICI combination regimens).

These recent data indicating that S RCC tumors are highly responsive to ICI have generated interest in determining the underpinnings of this responsiveness. Prior studies had suggested that S RCC tumors had increased tumor PD-L1 expression^41, 42^ and infiltration by CD8+ T cells^42^. These findings contrasted with another study that had reported that TGFβ signaling, which has been associated with immune exclusion and resistance to ICIs^43, 44^, was significantly increased in S RCC tumors^15^. More recently, two papers found that *CD274* (or PD-L1) gene amplifications are present in S RCC tumors and suggested that this genomic alteration may be underlying the increased PD-L1 tumor expression in these tumors and hypothesized that this genomic amplification may be underlying the immune responsiveness of S RCC tumors^2, 17^. In the present study, the integrative analysis of WES, RNA-seq, tumor PD-L1 expression by IHC, tumor CD8+ T cell infiltration by IF, and clinical outcomes on ICI monotherapy from pre-treatment samples of patients with metastatic renal cell carcinoma on two clinical trials (CheckMate 010 and CheckMate 025) allowed the in-depth examination of the immune characteristics of these tumors. The present study corroborated the finding of increased PD-L1 tumor cell expression in S/R RCC and found that CD8+ T cell infiltration tended to be increased in these tumors. We did not find *CD274* gene focal amplification to be enriched in these tumors compared to non-S/R RCC tumors. The small number of S/R RCC tumors that harbored *CD274* gene amplification and had PD-L1 expression data available all expressed tumor cell PD-L1. However, the increased expression of tumor cell PD-L1 in S/R RCC tumors and the responsiveness of these tumors to PD-1 inhibitor monotherapy appeared to be independent of *CD274* gene amplification (Fig 4c and Fig S10a-c). In addition, the analysis of two independent cohorts of RCC with RNA-seq (CheckMate and TCGA), revealed multiple previously unreported characteristics of the immune contexture of these tumors. First, all 8 “Hallmark” immune gene sets (but not the “Hallmark” TGFβ gene set), including IL6-JAK-STAT3 signaling and interferon gamma response, were enriched in S/R RCC tumors. Second, immune deconvolution revealed that multiple immune subsets that have previously been associated with an immune responsive microenvironment are significantly increased in S/R RCC tumors, including M1 macrophages, activated NK cells, and the Th1 T cell subset. These findings were also found to be largely consistent across S and R RCC subsets (Fig S8-9). Third, the expression of antigen presentation machinery genes, which has been found to correlate with increased cytotoxic immune infiltration and ICI responsiveness^23^, were significantly increased in S/R RCC tumors (Tables S5 and S7).

In order to evaluate whether sarcomatoid cell line models recapitulate the biology of S/R RCC tumors, we compared the transcriptional profiles of 6 sarcomatoid cell lines to 9 non-sarcomatoid cell lines. Although less statistically powered to detect similar effect sizes to those observed in the bulk tumor S/R vs. non-S/R RCC comparison (due to a smaller sample size), the transcriptional programs of these cell lines partially recapitulated the biology of S/R RCC tumors. In particular, EMT and apoptosis-caspase pathway gene sets were significantly enriched in both S/R RCC tumors and sarcomatoid cell lines. These results suggest that at least some of the transcriptional findings reported in this study for S/R RCC are driven by the sarcomatoid tumor cells themselves and that sarcomatoid cell lines could serve as adequate models for these tumors in future therapeutic development efforts for this RCC subtype. Since the transcriptional programs of cell lines have been suggested to be most predictive of their sensitivity profiles (as opposed to other molecular features)^34, 45^, these two signatures were then projected into two independent cell line drug screen datasets (CTRP and PRISM)^34, 46^. Sensitivity to CDK inhibitors appeared to correlate strongly with EMT and apoptosis-caspase pathway signatures in both datasets independently (Fig. S12-13 and Table S11). The CDK inhibitors that scored in these analyses target multiple CDKs, including those involved in transcription and cell cycle progression. We tested two CDKi (SNS-032 and alvocidib) along with a tyrosine kinase inhibitor control (axitinib) in two sarcomatoid and three non-sarcomatoid cell lines. The two sarcomatoid cell lines displayed decreased sensitivity to axitinib (a VEGF pathway inhibitor) as compared with the non-sarcomatoid cell line with the lowest EMT ssGSEA score, KMRC-20 (Fig. S12b and S14c), underscoring the limited response to this inhibitor of this canonical clear cell RCC pathway^47^ in these sarcomatoid cell lines. Sarcomatoid and non-sarcomatoid RCC cell lines showed globally similar sensitivities to the two CDKis tested in our assay. The overall sensitivity of both sarcomatoid and non-sarcomatoid RCC lines to the two CDKis tested may be explained by the specificities of the particular drugs tested as well as the plasticity in EMT gene expression program, even among non-S/R RCCs, that may modulate sensitivity to this class of agents. Study of the precise molecular determinants of response to these and other classes of therapeutic agents in S/R RCC is a ripe area for future investigation.

A limitation of this study is the potential bias induced by the inherent heterogeneity of S/R RCC tumors. Foci of sarcomatoid and rhabdoid features can be present anywhere within RCC tumors. When these tumors are being evaluated by pathologists, these foci of S/R features can be missed and S/R RCC tumors could be mis-classified as non-S/R RCC. In this study, we reviewed the pathology reports and slides of tumors (Methods) to attempt to minimize such misclassifications. Moreover, any biases due to misclassification would be expected to decrease the power of this study to detect an effect, thereby potentially increasing the risk of false negative but not false positive findings. In addition to misclassification, intra-tumoral histological heterogeneity (sarcomatoid/rhabdoid vs epithelioid foci within the same S/R RCC tumor in a patient) could also be associated with intra-tumoral molecular heterogeneity. In this study, using data from the present study and previously published studies, we find that the intra-tumoral mutational heterogeneity of S/R RCC tumors seems to be largely similar to that of non-S/R RCC tumors. In accordance with prior studies^14^, we find that mutations in certain genes (in particular *TP53*) may be enriched in S/R components of S/R RCC tumors. However, our overall analysis results suggest that mutational differences between S/R and non-S/R RCC tumors are greater than intra-tumoral mutational differences within S/R RCC tumors. The drivers of intra-tumoral histological heterogeneity require further evaluation and could be further investigated using novel single cell (DNA and/or RNA) and spatial transcriptomic methods.

In conclusion, our findings suggest that sarcomatoid and rhabdoid renal cell carcinoma tumors have distinctive genomic and transcriptomic features that may account for their aggressive clinical behavior. We also established that these tumors have significantly improved clinical outcomes on immune checkpoint inhibitors, which may be accounted for by an immune-inflamed phenotype; itself driven in part by upregulation of antigen presentation machinery genes in S/R RCC. Finally, our results suggest that sarcomatoid cell lines recapitulate the transcriptional programs of S/R RCC tumors and could serve as reasonably faithful models for these tumors, fueling the engine for future therapeutic discovery in this aggressive subtype of RCC. Further work is needed to determine whether other solid tumors with similar histological dedifferentiation components exhibit comparable molecular and clinical characteristics.

## Methods

### Clinical Cohorts and Patient Samples

The comparative clinical outcomes on immune checkpoint inhibitors (ICI) of patients with metastatic sarcomatoid and rhabdoid (S/R) renal cell carcinoma (RCC) were derived from: (1) CheckMate cohort (S/R RCC N = 120): two clinical trials evaluating an anti-PD-1 inhibitor (nivolumab) for metastatic clear cell RCC, CheckMate-025^27^ (NCT01668784) and CheckMate-010^26^ (NCT01354431), (2) Harvard cohort (S/R RCC N = 203): a retrospective cohort from the Dana-Farber/Harvard Cancer Center including patients from Dana-Farber Cancer Institute, Beth Israel Deaconess Medical Center, and Massachusetts General Hospital, (3) IMDC cohort (S/R RCC N = 927): a retrospective multi-center cohort of metastatic RCC that includes more than 40 international cancer centers and more than 10.000 patients with metastatic RCC. All patients had consented to an institutional review board (IRB) approved protocol to participate in the respective clinical trials and to have their samples collected for tumor and germline sequencing (for the CheckMate cohort) or to have their clinical data retrospectively collected for research purposes (Harvard and IMDC cohorts). Analysis was performed under a secondary use protocol, approved by the Dana-Farber Cancer Institute IRB. For all cohorts, the definition of sarcomatoid and rhabdoid RCC tumors was based on the ISUP 2013 consensus definitions: tumors were classified as harboring sarcomatoid features if they had any percentage of sarcomatoid component and as harboring rhabdoid features if they had any percentage of rhabdoid component (regardless of the background histology)^48^. For the Harvard and IMDC cohorts, sarcomatoid and rhabdoid status were determined by retrospective reviews of pathology reports. For the CheckMate cohort, sarcomatoid and rhabdoid features were retrospectively identified by review of pathology reports and of pathology slides by a pathologist. For the TCGA cohort, all pathology reports were first reviewed. Candidate sarcomatoid and/or rhabdoid cases were then reviewed by a pathologist. Cases that were unequivocal by the ISUP 2013 consensus definitions by pathology report and/or slide review were included. The TCGA cohort also included a subset of sarcomatoid RCC patients that had been previously retrospectively identified^15^. All pathology slides and reports for TCGA were accessed using cbioportal (https://www.cbioportal.org). Specifically, the following datasets were used: Kidney Renal Clear Cell Renal Cell Carcinoma (TCGA, Provisional), Kidney Chromophobe (TCGA, Provisional), Kidney Renal Papillary Cell Carcinoma (TCGA, Provisional). The sarcomatoid and rhabdoid annotations for the samples identified in TCGA are reported in Table S12. The clinical characteristics of the patients in the CheckMate cohort with molecular sequencing data were similar to those of the overall trial (Braun et al., *Nature Medicine*, in press).

### Cell Lines

Fifteen cell lines were acquired by our laboratory for baseline RNA-seq characterization including 6 that had been derived from sarcomatoid kidney cancer tumors (RCJ41M, RCJ41T1, RCJ41T2, BFTC-909, UOK127, and UOK276) and 9 that had been derived from non-sarcomatoid kidney cancer tumors (786-O, A498, ACHN, Caki-1, Caki-2, KMRC-1, KMRC-2, KMRC-20, and VMRC-RCZ). UOK127 and UOK276 were obtained from Dr. Linehan’s laboratory at the National Cancer Institute (NCI) while RCJ41M, RCJ41T1, and RCJ41T2 were obtained from Dr. Ho’s laboratory (Mayo Clinic, Phoenix, Arizona)^49^. Caki-1, Caki-2, A498, ACHN and 786-O were acquired from the American Type Culture Collection (ATCC). KMRC-1, KMRC-2, KMRC-20, VMRC-RCZ were obtained from JCRBbCell Bank and Sekisui XenoTech, LLC. BFTC-909 was obtained from Leibniz-Institut (DSMZ-Deutsche Sammlung von, Mikroorganismen und Zellkulturen GmbH).

Cell lines ACHN, VMRC-RCZ and 786-O were maintained in RPMI 1640 media (Gibco), supplemented with 10% FBS (Gibco) and 1% penicillin-streptomycin. Cell line A498 was maintained in EMEM media (Gibco), supplemented with 10% FBS (Gibco) and 1% penicillin-streptomycin. Caki-1 and Caki-2 were maintained in McCoy’s 5A media (Gibco), supplemented with 10% FBS (Gibco) and 1% penicillin-streptomycin. KMRC-1, KMRC-2, KMRC-20, UOK127, UOK276, BFTC-909, RCJ41T1, RCJ41T2 and RCJ41M were maintained in DMEM media (Gibco), supplemented with 10% FBS (Gibco) and 1% penicillin-streptomycin. Cultures were grown in a 37 °C incubator with 5% CO2. Total RNAs were isolated using the Trizol® reagent (Invitrogen), according to the manufacturer’s instructions.

For cell viability assays, cells were seeded in 96-well plates at densities ranging from 1,000-10,000 cells per well, depending on the cell line. After 24 hours, axitinib (S1005, Selleck), alvocidib (S1230, Selleck), or SNS-032 (S1145, Selleck) was added to cells at the indicated final concentrations. DMSO treatment was used as a negative control. Cell viability for 3 biological replicates of each treatment condition was assessed after 72 hours after drug treatment using the CellTiter-Glo Luminescent Cell Viability Assay (G7571, Promega) and an EnVision Multilabel Plate Reader (PerkinElmer). Viability was calculated for each cell line relative to its respective DMSO control wells.

### RNA and DNA Extraction, Sequencing, and Pre-processing

The methods used for DNA and RNA extraction and sequencing in the CheckMate 010 and 025 trials are described in a separate paper in more detail (Braun et al., *Nature Medicine*, in press). Briefly, archived formalin-fixed paraffin embedded (FFPE) tissue from pre-treatment samples of patients enrolled in these two trials were used. DNA and RNA were extracted from tumor samples along with paired germline DNA from whole blood. Germline and tumor DNA were sequenced using Illumina HiSeq2500 following a 2×100 paired-end sequencing recipe and targeting a depth of coverage of 100x. RNA was sequenced using a stranded protocol using Illumina HiSeq2500 following a 2×50 paired-end sequencing recipe and targeting a depth of 50 million reads. Mean exome-wide coverage for tumor samples was 129x and 112x for matched germline. For the RNA-seq data, the mean mapping rate of the included samples was 96.7% and mean number of genes detected was 21078.

For the TCGA cohort, publicly available data was downloaded for mutation data (https://gdc.cancer.gov/about-data/publications/mc3-2017), CNA data (https://www.cbioportal.org/datasets), upper-quartile (UQ) normalized transcripts-per-million (TPM) RNA-seq data (https://www.cbioportal.org/datasets), and clinical data (https://www.cbioportal.org/datasets)^50,51^. The dataset from the study by Malouf et al.^19^ of paired sequencing of sarcomatoid RCC was downloaded from https://www.nature.com/articles/s41598-020-57534-5#Sec16 (supplementary dataset 1). The dataset from the TRACERx Renal study^18^ was downloaded from https://www.ncbi.nlm.nih.gov/pmc/articles/PMC5938372/ (Tables S1 and S2).

For the OncoPanel cohort, DNA extraction and sequencing were performed as previously described for the OncoPanel gene panel assay^52^. The OncoPanel assay is an institutional analytic platform that is certified for clinical use and patient reporting under the Clinical Laboratory Improvement Amendments (CLIA) Act. The panel includes 275 to 447 cancer genes (versions 1 to 3 of the panel), including 239 genes that are common across all 3 versions of the panel. Mean sample-level coverage for the Oncopanel cohort was 305x.

For the 15 cell lines acquired by our laboratory, RNA-seq was done using Illumina Platform PE150 polyadenylated non-stranded sequencing. The average mapping rate was 98.9% and 17998 genes were detected on average (all RNA-seqQC2 quality control metrics are reported in Table S11). RNA-seq data (which were UQ normalized to an upper quartile of 1000 and log2- transformed) for 20 kidney cancer cell lines with RNA-seq and drug sensitivity data were downloaded from The Cancer Dependency Map Portal (DepMap)^53^ (https://depmap.org/portal/download/) and drug sensitivity data were downloaded from the Cancer Therapeutics Response Portal (CTRP v2)^46^ (https://portals.broadinstitute.org/ctrp/?cluster=true?page=#ctd2Cluster) and the PRISM 19Q4 secondary screen (https://depmap.org/portal/download/) as areas under the curve (AUC) for all agents.

### Genomic Analysis

The analytical pipeline for the WES data for the CheckMate 010 and 025 trials is described in detail in a separate paper (Braun et al., *Nature Medicine*, in press). Briefly, paired-end Ilumina reads were aligned to the hg19 human genome reference using the Picard pipeline (https://software.broadinstitute.org/gatk/documentation/tooldocs/4.0.1.0/ picard_fingerprint_CrosscheckFingerprints.php). Cross-sample contamination were assessed with the ContEst tool^54^, and samples with ≥5% contamination were excluded. Point mutations and indels were identified using MuTect^55^ and Strelka^56^, respectively. Possible artifacts due to orientation bias, germline variants, sequencing and poor mapping were filtered using a variety of tools including Orientation Bias Filter^57^, MAFPoNFilter^58^, and RealignmentFilter. Copy number events were called and filtered using GATK4 ModelSegments^59^. Copy number panel-of-normals was created based on matched germline samples. GISTIC^60^ was used to determine gene-level copy number alteration events. Clonality assessment was performed using ABSOLUTE^61^. Mutations were considered clonal if the expected cancer cell fraction (CCF) of the mutation as estimated by ABSOLUTE was 1, or if the estimated probability of the mutation being clonal was greater than 0.5. The intratumor heterogeneity index (ITH) was defined as the ratio of subclonal mutations to clonal mutations.

OncoPanel mutation and gene-level copy number calling was performed as previously described^52^. In particular, variants were filtered to exclude those that occurred at a frequency of >0.1% in the Exome Sequencing Project database (http://evs.gs.washington.edu/EVS/) in order to remove variants that were probably germline variants. Additionally, in order to further remove potential germline variants from the OncoPanel results, Ensembl Variant Effect Predictor (VEP)^62^ was run on the OncoPanel mutations and mutations present at an allelic frequency of 0.5% in one of the superpopulations were excluded from all downstream analyses.

For the purposes of the present genomic analysis, mutation and CNA of 244 genes were analyzed (Table S13), including the 239 genes that are common across the 3 versions of the panel, 3 frequently mutated genes in RCC (*KDM5C*, *KMT2D*, and *PBRM1*)^16^ that are only included in versions 2 and 3 of the panel, and 2 genes that are included in none of the 3 versions of the panel, including a frequently mutated RCC gene (*KMT2C*)^16^ and a gene that has been previously suggested to be more frequently mutated in sarcomatoid RCC (*RELN*)^15^. All mutations from TCGA, Oncopanel, and CheckMate cohorts were annotated using Oncotator^63^ (https://software.broadinstitute.org/cancer/cga/oncotator). For WES data, only mutations with more than 30x coverage were included.

Somatic genomic alterations (mutations and insertions-deletions [indels]) were considered to be pathogenic if they were truncating (nonsense or splice site), indels, or missense mutations that were predicted to be pathogenic by Polyphen-2 HumDiv score^64^ ≥0.957 or Mutation Assessor^65^ score >1.90. Tumor mutational burden was calculated as the sum of all non-synonymous mutations divided by the estimated bait set (30 Megabases [Mb] for WES, 1.32 Mb for panel v3, 0.83 Mb for panel v2, and 0.76 Mb for panel v1). Moreover, the indel burden (either all indels or only frameshift indels) was normalized by dividing by the estimated bait set for each version of OncoPanel. Gene-level deep deletions and high amplifications were considered for the primary copy number analysis, while any deletions (one-copy or two-copy) and any amplifications (low or high) were analyzed as a supplementary analysis.

The co-mutation plot was generated excluding patients that had either mutation or CNA data missing in any of the 3 cohorts (as reported in Table S1). The estimate of percentage mutated took into account the missing genes for patients sequenced by panel sequencing (these percentages were estimated while excluding patients sequenced by panel sequencing for *RELN* and *KMT2C*, while only the patients sequenced by panel v1 were excluded for *KDM5C*, *KMT2D*, and *PBRM1).* TMB was compared between S/R and non-S/R in each of the three cohorts independently using Mann-Whitney U tests. Genomic alterations (mutations and indels, deep deletions, and high amplifications analyzed separately) were compared between S/R and non-S/R in each of the three cohorts independently using a Fisher’s exact test. For the OncoPanel cohort, for *KDM5C*, *KMT2D*, and *PBRM1*, patients that had been sequenced by panel version 1 were excluded from the analysis. Only genes that were altered in at least 5% of patients (in all patients with RCC or in the S/R RCC group) in at least one of the 3 cohorts were tested. The p-values from the 3 cohorts were subsequently combined using Fisher’s method for meta-analyses. The combined p-values were corrected for multiple hypothesis testing using Benjamini-Hochberg correction. Findings were considered to be significant if they were statistically significant at q<0.05 and the same direction of the effect was observed in at least two of the three included datasets.

For the analysis of paired data in the dataset by Malouf et al. (paired sarcomatoid and epithelioid regions of S RCC tumors), continuous variables were compared by the paired Wilcoxon signed rank test. Mutation rates in genes were compared using McNemar’s test.

### Transcriptomic Analysis

RNA-seq data from the CheckMate cohorts and the 15 cell lines sequenced in our laboratory were aligned using STAR^66^, quantified using RSEM^67^, and evaluated for quality using RNA-seqQC2^68^. Samples were excluded if they had an interquartile range of log2(TPM+1)<0.5 or had less than 15,000 genes detected. Additionally, since the CheckMate cohort had been sequenced by a stranded protocol, samples were filtered if they had an End 2 Sense Rate<0.90 or End 1 Sense Rate>0.10 (as defined by RNA-seqQC2). For samples where RNA-seq was performed in duplicates, the run with a higher interquartile range of log2(TPM+1), considered a surrogate for better quality data, was used. We subsequently filtered genes that were not expressed in any of the samples (in each cohort independently) then UQ-normalized the TPMs to an upper quartile of 1000, and log2-transformed them. Since the CheckMate cohort had been sequenced in 4 separate batches, principal component analysis (PCA) was used to evaluate for batch effects and 4 batches were observed. These 4 batches were corrected for using ComBat^69^ (Fig. S15). Subsequently, a PCA was performed on the ComBat-corrected expression matrix to confirm that batch effects had been adequately corrected for (Fig. S15). Moreover, a constant that was equal to the first integer above the minimum negative expression value obtained post-ComBat (constant of +21) was added to eliminate negative gene expression values that were a by-product of ComBat correction. The ComBat-corrected expression matrix was used for all downstream analyses on the CheckMate cohort. All downstream analyses were computed on the TCGA and CheckMate cohorts independently and only results which were found to be independently statistically significant in each of the two cohorts were considered to be significant.

GSEA between S/R and non-S/R was run using the Java Application for GSEA v4.0.0 and MSigDB 7.0^70^ on the 50 “Hallmark” gene sets, *MYC* v1 and v2 “Founder” gene sets, and select KEGG^25^ and REACTOME^24^ antigen presentation machinery gene sets. Gene sets were considered to be enriched if q<0.25. Single sample GSEA (ssGSEA) was additionally computed using the “GSVA” package^71^ in the R programming environment to obtain sample-level GSEA scores. Differential gene expression analysis was computed using the non-parametric Mann-Whitney U test and Benjamini-Hochberg false discovery rate correction with q<0.05 considered statistically significant. The CIBERSORTx deconvolution algorithm^72^ was used to infer immune cell infiltration from RNA-seq data (Job type: “Impute cell fractions”), in absolute mode, on the LM22 signature^73^, with B mode batch correction (in order to correct for the batch effect between the LM22 signature, which was derived from microarray data, and the data used in this study which consisted of RNA-seq), with quantile normalization disabled, and in 1000 permutations. All samples which had a p-value for deconvolution >0.05 were considered to have failed deconvolution and were therefore discarded from all downstream analyses. Relative cell proportions were obtained by normalizing the CIBERSORTx output to the sample-level sum of cell counts (in order to obtain percentages of immune infiltration). A constant of 10^- 06 was added to all proportions in order to allow the computation of immune cell ratios. Additionally, Th1, Th2, and Th17 scores were computed using ssGSEA (and were normalized to scores between 0 and 100) based on previously described signatures for these cell types^30^. All immune cell proportions and ratios were compared between S/R and non-S/R using a non-parametric Mann-Whitney U test with Benjamini-Hochberg correction and a q-value threshold of 0.05 for statistical significance.

In order to evaluate whether specific signatures predicted outcomes in S/R RCC, Cox regression models were performed to evaluate the relationship between ssGSEA scores, modeled as continuous variables (multiplied by a factor of 100), and survival outcomes. ssGSEA scores found to be significantly associated with survival outcomes were used to dichotomize S/R RCC patients into two groups at the median of the score. The dichotomized groups were evaluated using Kaplan-Meier curves and compared using log-rank tests. In order to evaluate whether such relationships held in patients with non-S/R RCC, the same analysis was conducted in non-S/R RCC using the ssGSEA scores that were found to be related to outcomes in S/R RCC. In addition, for non-S/R RCC patients, the group was also dichotomized based on the median of the S/R RCC group and compared by Kaplan-Meier methodology and log-rank tests. In particular, this was done for MYC v1 scores which were found to be significantly related to outcomes in the S/R RCC group and not found to be related to outcomes when evaluated continuously in the non-S/R RCC group or when dichotomized at the median.

### Cell Line In Silico Drug Sensitivity Analysis

In order to evaluate potential novel therapeutic targets for S/R RCC, we computed ssGSEA scores for the 20 kidney cancer cell lines in DepMap that also had drug sensitivity data reported as areas under the curve (AUCs) of the dose-response curve in CTRP v2 and in the PRISM secondary screen. Using the gene signatures that were found to be significantly upregulated in both bulk tumor RNA-seq cohorts (in the TCGA and CheckMate cohorts independently) and sarcomatoid cell lines, we correlated the scores to drug sensitivity AUC data using Pearson’s r correlation coefficients. Only therapeutic agents that were tested in at least 8 of the 20 kidney cancer cell lines were evaluated in CTRP v2. For visualization, the ssGSEA-AUC correlations were grouped by drug types and illustrated in a heatmap (in which negative correlations indicated that higher ssGSEA scores correlated with lower AUCs and therefore greater sensitivity). Moreover, scatter plots of the correlations were displayed for key correlations.

### Immunohistochemistry and Immunofluorescence

PD-L1 expression on the membrane of tumor cells was assessed using the Dako assay, as previously described in the CheckMate 025 and 010 trials^26, 27^. Tumors were considered PD-L1 positive if they expressed PD-L1 on ≥1% of tumor cells.

The immunofluorescence assay used is described in detail in a separate paper (Braun et al., *Nature Medicine*, in press). CD8 immunostain was performed as part of a multiplex fluorescent IHC panel on 4 μm FFPE sections. Tumor sections were stained using the Opal multiplex IHC system (PerkinElmer), which is based on tyramide-conjugated fluorophores. All slides were counterstained with Spectral DAPI (PerkinElmer) and manually coverslipped. The slides were imaged using the Vectra 3 automated quantitative pathology imaging system (PerkinElmer) and whole slide multispectral images were acquired at 10x magnification.

Digital whole slide multispectral images were then uploaded into HALO Image Analysis platform version 2.1.1637.18 (Indica Labs). For each case, the tumor margin and center were defined while also excluding empty spaces, necrosis, red blood cells and fibrotic septa. Specifically, the tumor margin was defined as the space within 500 μm (in either direction) of the interface between the tumor and surrounding tissue. Image analysis algorithms were built using Indica Labs High-Plex FL v2.0 module to measure the area within each layer, perform DAPI-based nuclear segmentation and detect CD8 (FITC)-positive cells by setting a dye cytoplasm positive threshold. A unique algorithm was created for each tumor and its accuracy was validated through visual inspection by at least one pathologist.

### Clinical Outcomes

For patients in the Harvard and IMDC cohorts, clinical data were retrospectively collected. OS was defined as the time from the start of the line of therapy (ICI or non-ICI) until death from any cause. Time to treatment failure (TTF) was defined as the time from start of the line of therapy until discontinuation of therapy for any cause. Since assessment of responses in these retrospective cohorts was not subject to radiological review specifically for the purpose of this study, responses were defined based on RECIST v1.1 criteria^74^ as available by retrospective review. For the CheckMate cohort, OS was defined from the time of randomization until death from any cause. Progression free survival (PFS) was defined from randomization until death or progression. Both PFS and ORR were defined using RECIST v1.1 criteria. All patients who were lost to follow-up or did not have an event at last follow-up were censored.

### Statistical Analysis

The dose-response curves for the in vitro cell viability assays performed at DFCI were generated using GraphPad PRISM 8. All analyses were done in the R programming environment version 3.6.1. For boxplots, the upper and lower hinges represent the 75^th^ and 25^th^ percentiles, respectively. The whiskers extend in both directions until the largest or lowest value not further than 1.5 times the interquartile range from the corresponding hinge. Outliers (beyond 1.5 times the interquartile range) are plotted individually. Continuous variables were summarized by their means and standard deviations (SD) or medians and interquartile ranges (IQR) or ranges. Categorical variables (such as gene alterations) were summarized by their percentages. For survival outcomes, the Kaplan-Meier methodology was used to summarize survival distributions in different groups; 18-month PFS (or TTF) and 2-year OS were provided with 95% confidence intervals. For survival outcomes, multivariable Cox regression models were used for the comparison of ICI and non-ICI regimens and adjusted hazard ratios (HR) with their 95% confidence intervals were reported. Specifically, the IMDC risk groups^75^ (Poor vs. Intermediate/Favourable), line of therapy (2nd line and beyond vs. 1st line), and background histology (clear cell vs. non-clear cell) were adjusted for in the Harvard and IMDC cohort analyses and the Memorial Sloan Kettering Cancer Center (MSKCC) risk groups^76^ (Poor vs. Intermediate vs. Favourable) were adjusted for in the CheckMate cohort analysis. Similarly, the ORR was compared between the ICI and non-ICI using multivariable logistic regression models adjusting for the same covariates (except for the CheckMate cohort, in which only one patient had had a response in the everolimus arm and therefore the adjusted odds ratio was not estimable). For all multivariable analyses, patients with missing data in any of the variables were excluded from the analysis. For ORR analyses, only patients who were evaluable for response were included in the analysis. The Kaplan-Meier methodology for assessing point estimates of survival was computed using the “landest” package in R. All heatmaps were created using the R package “pheatmap” and were computed based on Z score transformations. When multiple cohorts were represented in the same heatmap, the Z score normalization was done within each cohort separately (in order to account for batch effects in visualization). All tests were two-tailed and considered statistically significant for p<0.05 or q<0.05 unless otherwise specified.

## Data Availability

All relevant correlative data are available from the authors and/or are included with the manuscript. All clinical and correlative data from the CheckMate 010 and 025 clinical trials are made separately available as part of the accompanying paper (Braun et al., *Nature Medicine*, in press). All intermediate data from the RNA-seq analyses of the CheckMate and TCGA cohorts are made available in tables S6 (single sample gene set enrichment analysis scores) and S9 (CIBERSORTx immune deconvolution). The raw, transformed, and intermediate data from the generated cell line RNA-seq data are made available in Table S11. Any other queries about the data used in this study should be directed to the corresponding authors of this study.

## Code Availability

Algorithms used for data analysis are all publicly available from the indicated references in the Methods section. Any other queries about the custom code used in this study should be directed to the corresponding authors of this study.

## Acknowledgements

We thank the OncoPanel study team and the patients who contributed their data to research and participated in clinical trials. We thank all contributors to The International Metastatic Renal-Cell Carcinoma Database Consortium for their data contributions. We thank Dr. Kanishka Sircar and Dr. Jose Antonio Karam for providing us with additional data and insights from their previously published study. This work was supported in part by Dana-Farber / Harvard Cancer Center Kidney Cancer SPORE (P50-CA101942-12), DOD CDMRP (W81XWH-18-1-0480), and Bristol-Myers Squibb. S.A.S acknowledges support by the NCI (R50RCA211482). RF has been funded in part by the ARC foundation grant. XXW has been funded in part by the KCA YIA. CJW is supported in part by The G. Harold and Leila Y. Mathers Foundation, and she is a Scholar of the Leukemia and Lymphoma Society. T.H.H is supported by National Cancer Institute (R01 CA224917). E.M.V is supported by NIH R01 CA227388. T.K.C. is supported in part by the Dana-Farber/Harvard Cancer Center Kidney SPORE and Program, the Kohlberg Chair at Harvard Medical School and the Trust Family, Michael Brigham, and Loker Pinard Funds for Kidney Cancer Research at DFCI.

## Author Contributions

Conception and Design: Ziad Bakouny, David A. Braun, Eliezer M. Van Allen, Toni K. Choueiri

Provision of study material or patients: Ziad Bakouny, Wenting Pan, Gwo-Shu Mary Lee, Stephen Tang, Kevin Bi, Jihye Park, Sabrina Camp, Maura Sticco-Ivins, Laure Hirsch, Giannicola Genovese, Michelle S. Hirsch, Srinivas Raghavan Viswanathan, Steven L. Chang, Xiao X. Wei, Bradley A. McGregor, Lauren C. Harshman, Leigh Ellis, Mark Pomerantz, Matthew L. Freedman, Michael B. Atkins, Catherine J. Wu, Thai H. Ho, W. Marston Linehan, David F. McDermott, Daniel Y.C. Heng, Sabina Signoretti, Eliezer M. Van Allen, and Toni K. Choueiri

Collection and assembly of data: Ziad Bakouny, Stephen Tang, Srinivas Raghavan Viswanathan, Xin Gao, Abdallah Flaifel, Amin H. Nassar, Sarah Abou Alaiwi, Megan Wind-Rotolo, Petra Ross-Macdonald, Ronan Flippot, Gabrielle Bouchard, John A. Steinharter, Pier Vitale Nuzzo, Miriam Ficial, Miriam Sant’Angelo, Jacob E. Berchuck, Shaan Dudani

Data Analysis and Interpretation: Ziad Bakouny, Stephen Tang, Srinivas Raghavan Viswanathan, David A. Braun, Sachet A. Shukla, Yue Hou, Wanling Xie, Megan Wind-Rotolo, Petra Ross-Macdonald, Alice Bosma-Moody, Meng Xiao He, Natalie Vokes, Jackson Nyman, Juliet Forman, Maxine Sun, Eliezer M. Van Allen, Toni K. Choueiri

Manuscript writing and revision: All authors

Final approval of manuscript: All authors

Accountable for all aspects of work: All authors

## Competing Interests statement

ZB: reported nonfinancial support from Bristol-Myers Squibb and Genentech/ImCore.

D.A.B. reported nonfinancial support from, Bristol-Myers Squibb, and personal fees from Octane Global, Defined Health, Dedham Group, Adept Field Solutions, Slingshot Insights, Blueprint Partnerships, Charles River Associates, Trinity Group, and Insight Strategy, outside of the submitted work.

S.A.S. reported nonfinancial support from Bristol-Myers Squibb, and equity in 152 Therapeutics outside the submitted work.

X.G: Research Support to Institution: Exelixis. X.X.W: Research Support: BMS.

B.A.M is a consultant for Bayer, Astellas, Astra Zeneca, Seattle Genetics, Exelixis, Nektar, Pfizer, Janssen, Genentech and EMD Serono. He received research support to Dana Farber Cancer Institute (DFCI) from Bristol Myers Squibb, Calithera, Exelixis, Seattle Genetics.

L.C.H reports consulting fees from Genentech, Dendreon, Pfizer, Medivation/Astellas, Exelixis, Bayer, Kew Group, Corvus, Merck, Novartis, Michael J Hennessy Associates (Healthcare Communications Company and several brands such as OncLive and PER), Jounce, EMD Serrano, and Ology Medical Education; Research funding from Bayer, Sotio, Bristol-Myers Squib, Merck, Takeda, Dendreon/Valient, Jannsen, Medivation/Astellas, Genentech, Pfizer, Endocyte (Novartis), and support for research travel from Bayer and Genentech.

M.B.A: Advisory Board participation: BMS, Merck, Novartis, Arrowhead, Pfizer, Galactone, Werewolf, Fathom, Pneuma, Leads BioPharma; Consultant: BMS, Merck, Novartis, Pfizer, Genentech-Roche, Exelixis, Eisai, Aveo, Array, AstraZeneca, Ideera, Aduro, ImmunoCore, Boehringer-Ingelheim, Iovance, Newlink, Pharma, Surface, Alexion, Acceleron, Cota, Amgen; Research Support to institution: BMS, Merck, Pfizer, Genentech.

C.J.W: Co-founder of Neon Therapeutics, and is a member of its SAB. Receives research funding from Pharmacyclics.

T.H.H: Advisory board participation: Surface Therapeutics, Exelixis, Genentech, Pfizer, Ipsen, Cardinal Health; research support-Novartis.

D.F.M reports a consulting/advisory role for Bristol-Myers Squibb, Merck, Roche/Genentech, Pfizer, Exelixis, Novartis, Eisai, X4 Pharmaceuticals, and Array BioPharma; and reports that his home institution receives research funding from Prometheus Laboratories.

D.H: consulting or research funding from Pfizer, Novartis, BMS, Ipsen, Exelixis, and Merck.

S.S: Research support to Institution: Bristol-Myers Squibb, AstraZeneca, Novartis, Exelixis; Consultant: Merck, AstraZeneca, Bristol-Myers Squibb, AACR, and NCI; royalties: Biogenex.

E.M.V: Advisory/Consulting: Tango Therapeutics, Genome Medical, Invitae, Illumina, Ervaxx; Research support: Novartis, BMS; Equity: Tango Therapeutics, Genome Medical, Syapse, Ervaxx, Microsoft; Travel reimbursement: Roche/Genentech; Patents: Institutional patents filed on ERCC2 mutations and chemotherapy response, chromatin mutations and immunotherapy response, and methods for clinical interpretation.

T.K.C: Research (Institutional and personal): AstraZeneca, Alexion, Bayer, Bristol Myers-Squibb/ER Squibb and sons LLC, Cerulean, Eisai, Foundation Medicine Inc., Exelixis, Ipsen, Tracon, Genentech, Roche, Roche Products Limited, F. Hoffmann-La Roche, GlaxoSmithKline, Lilly, Merck, Novartis, Peloton, Pfizer, Prometheus Labs, Corvus, Calithera, Analysis Group, Sanofi/Aventis, Takeda, National Cancer Institute (NCI), National Institute of Health (NIH), Department of Defense (DOD).; Honoraria: AstraZeneca, Alexion, Sanofi/Aventis, Bayer, Bristol Myers-Squibb/ER Squibb and sons LLC, Cerulean, Eisai, Foundation Medicine Inc., Exelixis, Genentech, Roche, Roche Products Limited, F. Hoffmann-La Roche, GlaxoSmithKline, Merck, Novartis, Peloton, Pfizer, EMD Serono, Prometheus Labs, Corvus, Ipsen, Up-to-Date, NCCN, Analysis Group, NCCN, Michael J. Hennessy (MJH) Associates, Inc (Healthcare Communications Company with several brands such as OnClive, PeerView and PER), Research to Practice, L-path, Kidney Cancer Journal, Clinical Care Options, Platform Q, Navinata Healthcare, Harborside Press, American Society of Medical Oncology, NEJM, Lancet Oncology, Heron Therapeutics, Lilly, ASCO, ESMO ; Consulting or Advisory Role: AstraZeneca, Alexion, Sanofi/Aventis, Bayer, Bristol Myers-Squibb/ER Squibb and sons LLC, Cerulean, Eisai, Foundation Medicine Inc., Exelixis, Genentech, Heron Therapeutics, Lilly, Roche, GlaxoSmithKline, Merck, Novartis, Peloton, Pfizer, EMD Serono, Prometheus Labs, Corvus, Ipsen, Up-to-Date, NCCN, Analysis Group, Pionyr, Tempest.; No speaker’s bureau; Stock ownership: Pionyr, Tempest.; No leadership or employment in for-profit companies. Other present or past leadership roles: Director of GU Oncology Division at Dana-Farber and past President of medical Staff at Dana-Farber), member of NCCN Kidney panel and the GU Steering Committee, past chairman of the Kidney Cancer Association Medical and Scientific Steering Committee); Patents, royalties or other intellectual properties: International Patent Application No. PCT/US2018/12209, entitled “PBRM1 Biomarkers Predictive of Anti-Immune Checkpoint Response,” filed January 3, 2018, claiming priority to U.S. Provisional Patent Application No. 62/445,094, filed January 11, 2017 and International Patent Application No. PCT/US2018/058430, entitled “Biomarkers of Clinical Response and Benefit to Immune Checkpoint Inhibitor Therapy,” filed October 31, 2018, claiming priority to U.S. Provisional Patent Application No. 62/581,175, filed November 3, 2017; Travel, accommodations, expenses, in relation to consulting, advisory roles, or honoraria; Medical writing and editorial assistance support may have been funded by Communications companies funded by pharmaceutical companies (ClinicalThinking, Envision Pharma Group, Fishawack Group of Companies, Health Interactions, Parexel, Oxford PharmaGenesis, and others); The institution (Dana-Farber Cancer Institute) may have received additional independent funding of drug companies or/and royalties potentially involved in research around the subject matter; CV provided upon request for scope of clinical practice and research; Mentored several non-US citizens on research projects with potential funding (in part) from non-US sources/Foreign Components: Asmar Wood S.A.L. is a private company based in Beirut, Lebanon that will provide a total of $100,000 in salary support to Dr. Sarah Abou Alaiwi from 7/1/2018 to 7/1/2020 during her post-doctoral research fellowship at DFCI; Fondation Arc Pour La Recherche Sur Le Cancer is a not-for-profit foundation based in Villejuif, France that provides 2561.04€ per month in salary support to Dr. Ronan Flippot during his clinical training at DFCI from 5/2/2018 to 11/4/2018.

The other authors declare no potential conflicts of interest.

## Supplementary Figure Legends

**Figure S1:**
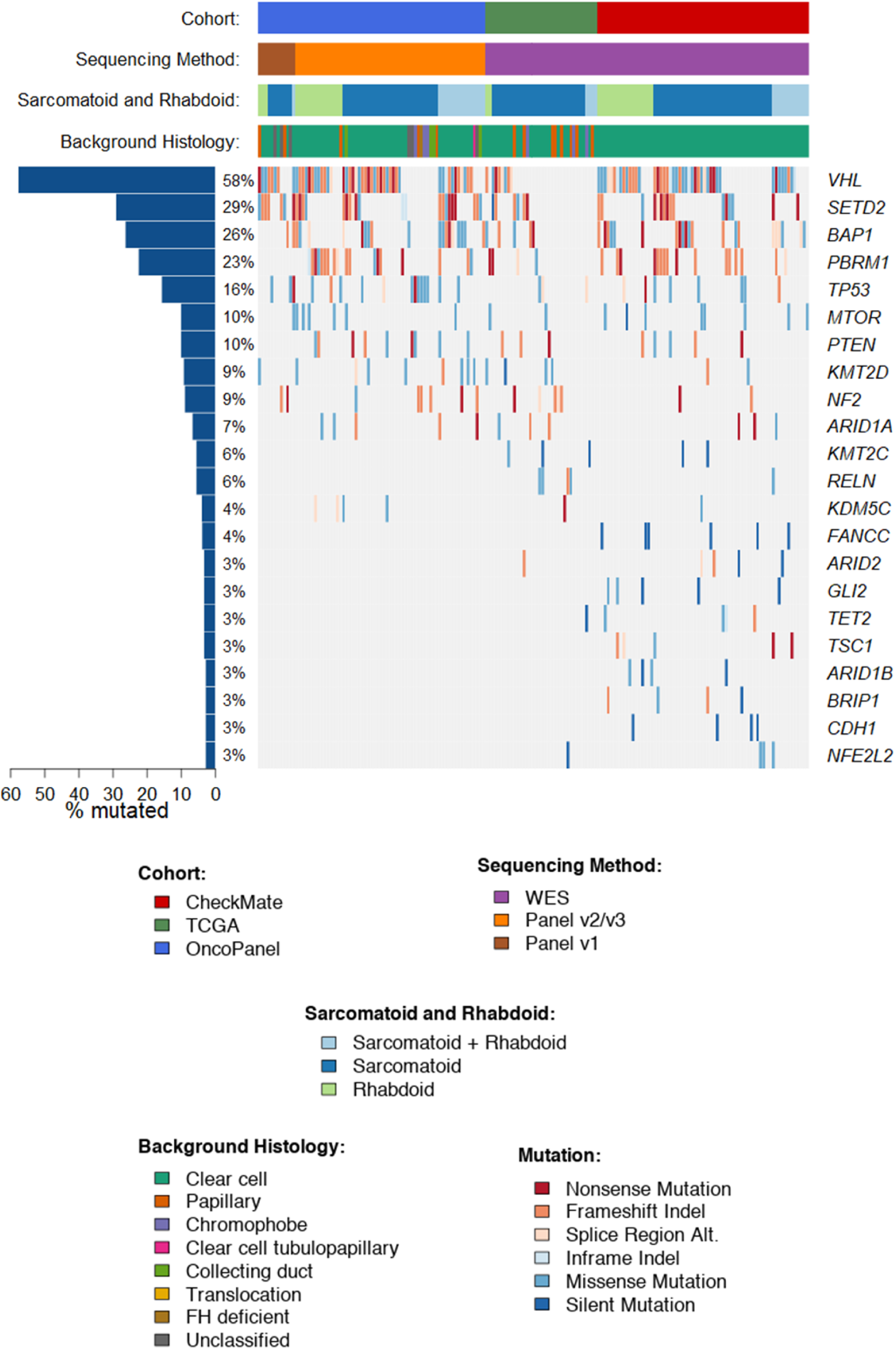
Co-mutation plot of patients with S/R RCC across the CheckMate, OncoPanel, and TCGA cohorts (in relation to Fig. 1). OncoPanel (all versions) did not include *KMT2C* or *RELN*. OncoPanel v1 did not include *KDM5C*, *KMT2D*, or *PBRM1* genes. The percentage mutated numbers take this into account by excluding the corresponding patients from the percentage calculation. Alt: Alteration; TCGA: The Cancer Genome Atlas; WES: Whole Exome Sequencing

**Figure S2:**
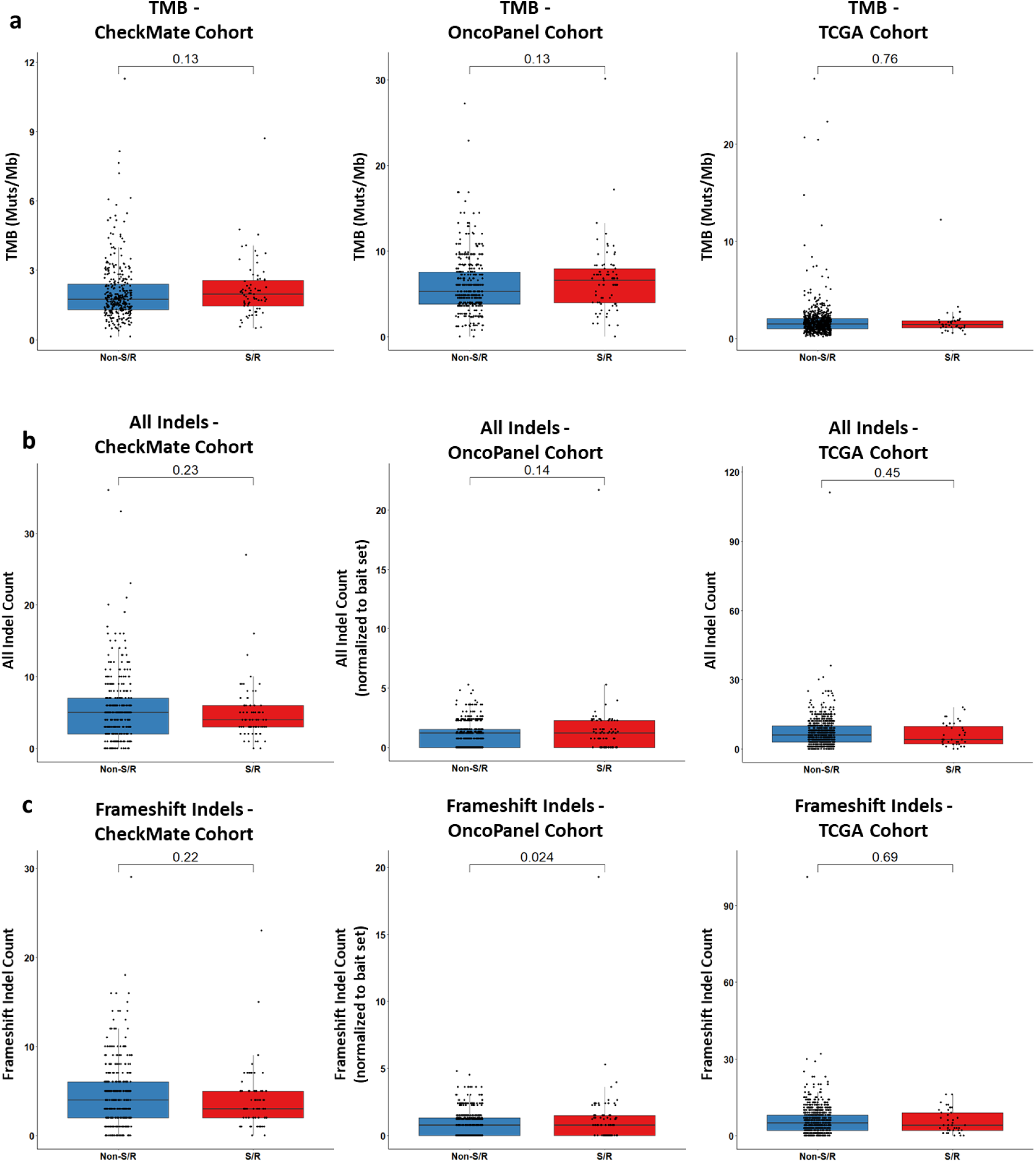
S/R RCC tumors have a similar overall (a) tumor mutational burden, (b) total indel load, and (c) frameshift indel load compared to non-S/R RCC tumors in the CheckMate, TCGA, and OncoPanel cohorts (in relation to Fig. 1). Mann-Whitney U test p-values shown. Muts: Mutations; Mb: Megabase; S/R: Sarcomatoid/Rhabdoid; TMB: Tumor Mutational Burden.

**Figure S3:**
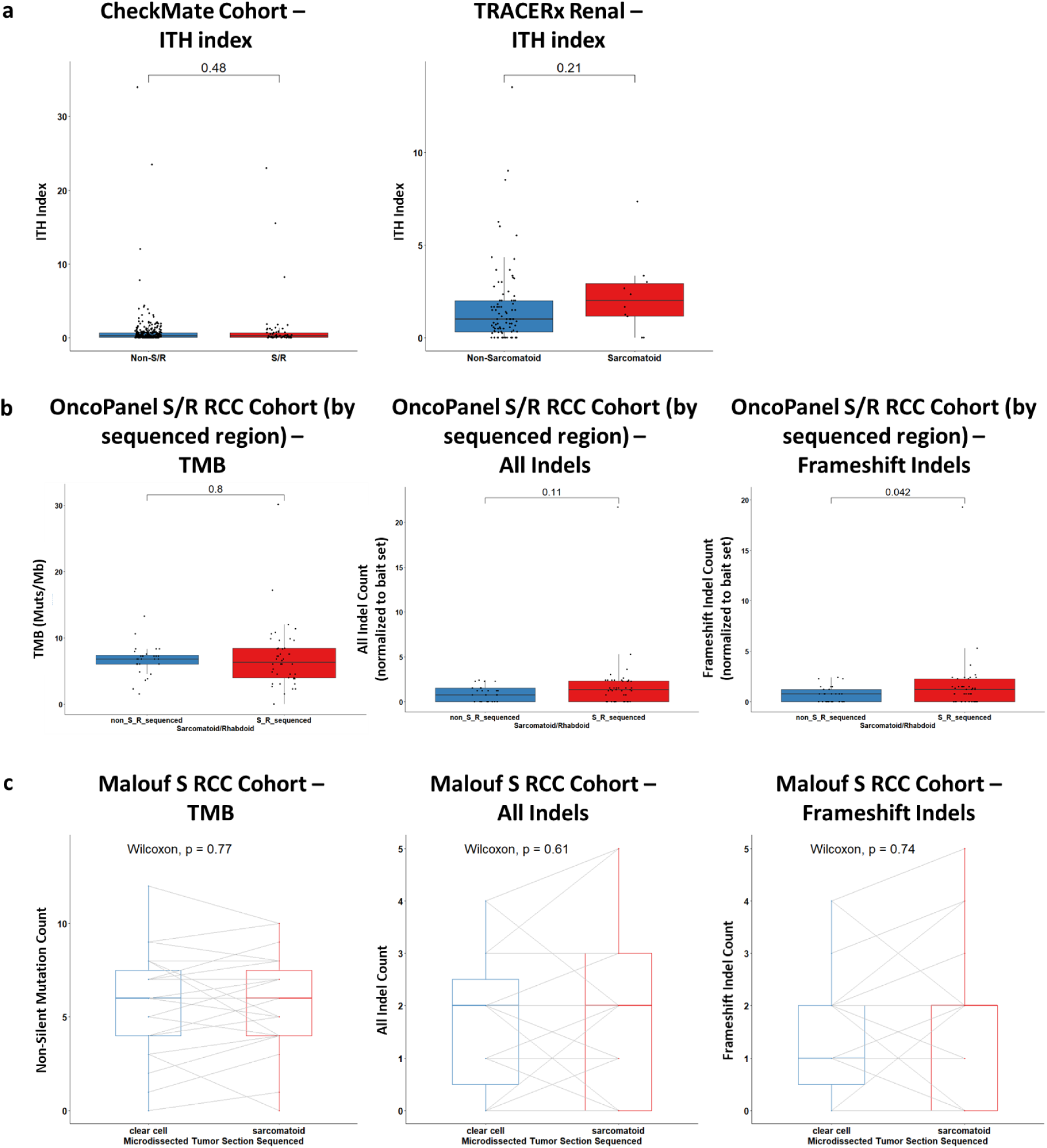
Limited intra-tumoral mutational heterogeneity of S/R RCC tumors (in relation to Fig. 1). (a) S/R RCC tumors have a similar intra-tumoral heterogeneity index to non-S/R RCC tumors in the CheckMate and TRACERx Renal cohorts. (b) Similar tumor mutational burden, total indel load, and frameshift indel load between the mesenchymal (S/R) and epithelioid (non-S/R) components within S/R RCC tumors in the OncoPanel cohort. (c) Similar tumor mutational burden, total indel load, and frameshift indel load between the mesenchymal (S) and epithelioid or clear cell (non-S) components within S RCC tumors in the Malouf cohort. Mann-Whitney U test p-values shown in (a) and (b). Paired Wilcoxon signed rank test p-value shown in (c). Muts: Mutations; Mb: Megabase; S/R: Sarcomatoid/Rhabdoid; TMB: Tumor Mutational Burden.

**Figure S4:**
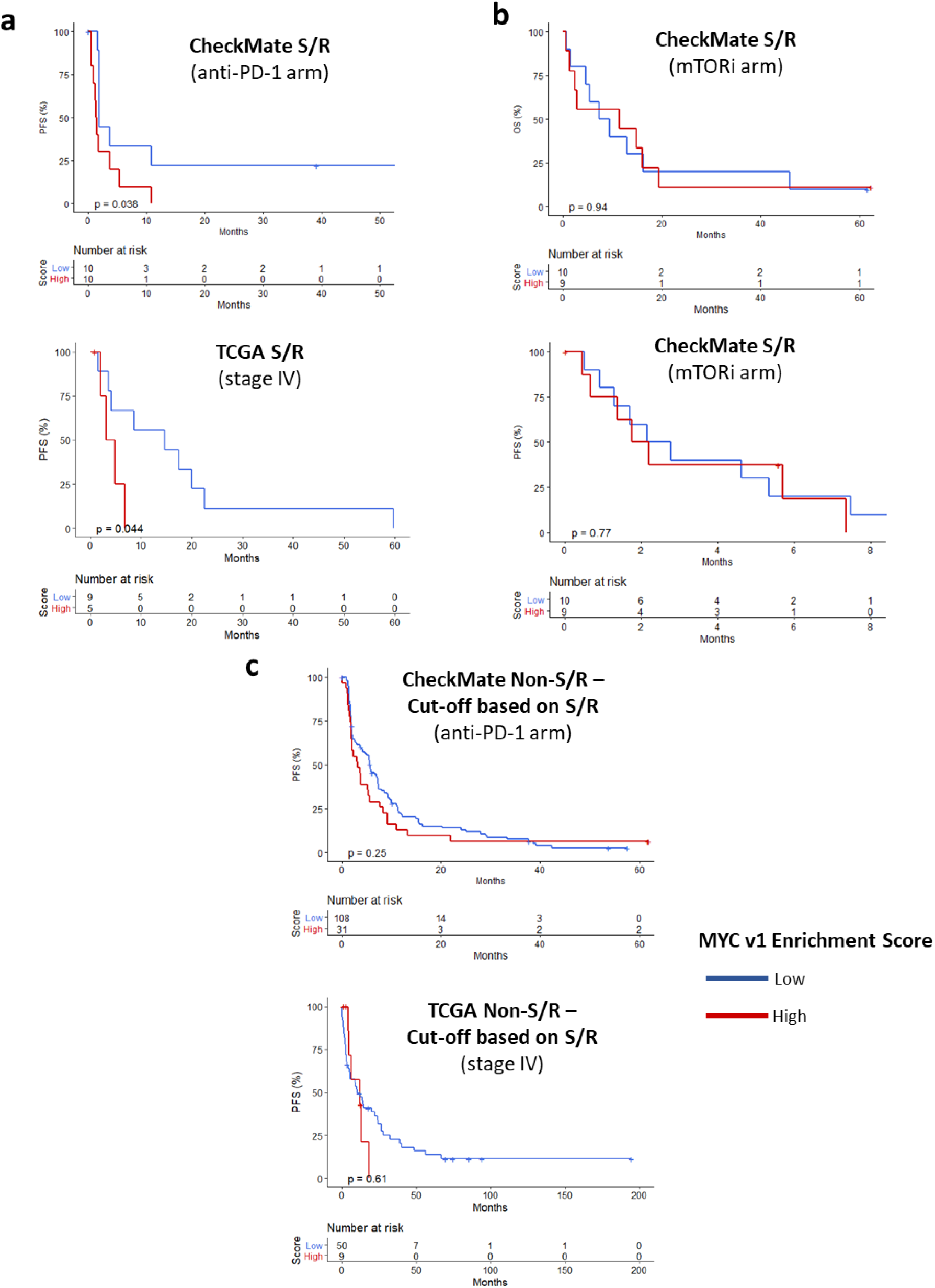
Transcriptional profiling of S/R RCC reveals the molecular correlates of its poor prognosis and identifies subsets of non-S/R tumors associated with a poor prognosis (in relation to Fig. 2). (a) Kaplan-Meier curves for PFS by *MYC* v1 score within the S/R group of the CheckMate (anti-PD-1 arm) and TCGA (stage IV) cohorts; *MYC* v1 score dichotomized at the median. (b) Kaplan-Meier curves for OS and PFS by *MYC* v1 score within the S/R group of the CheckMate (mTORi arm) cohort; *MYC* v1 score dichotomized at the median. (c) Kaplan-Meier curves for PFS by *MYC* v1 score within the non-S/R group of the CheckMate (anti-PD-1 arm) and TCGA (stage IV) cohorts; *MYC* v1 score dichotomized at the median of the S/R group. *MYC* v1: *MYC* Targets Version 1; S/R: Sarcomatoid/Rhabdoid; TCGA: The Cancer Genome Atlas; mTORi: Mammalian Target of Rapamycin Inhibitors

**Figure S5:**
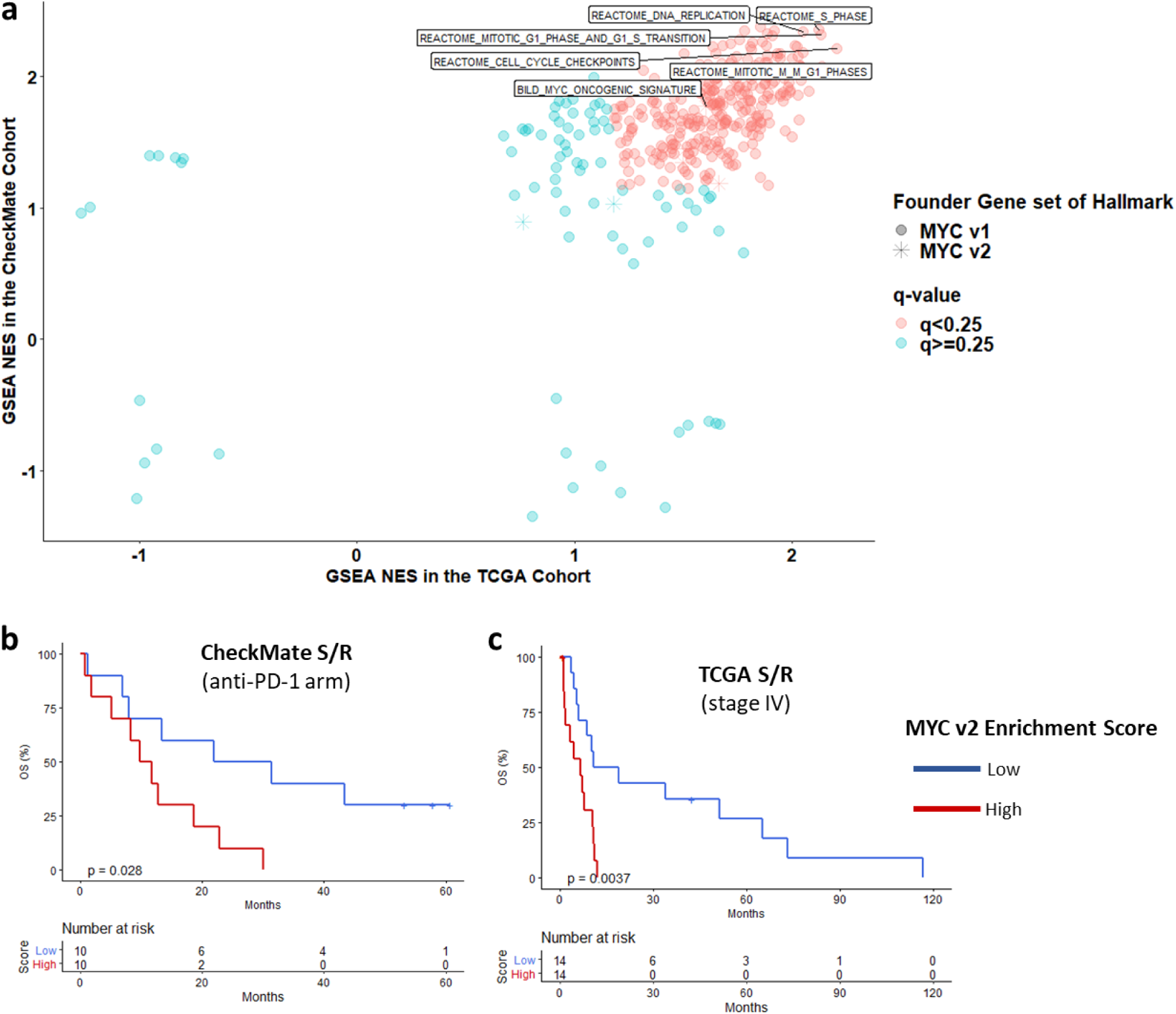
Upregulation of MYC-regulated gene expression and correlation with outcomes in S/R RCC (in relation to Fig. 2). (a) Enrichment of “Founder” gene sets of the “Hallmark” *MYC* v1 and v2 gene sets in the CheckMate and TCGA cohorts by GSEA. Kaplan-Meier curves for OS by *MYC* v2 score within the S/R group of the CheckMate (anti-PD-1 arm) and TCGA (stage IV) cohorts; *MYC* v1 score dichotomized at the median. GSEA: Gene Set Enrichment Analysis; *MYC* v2: *MYC* Targets Version 2; NES: Normalized Enrichment Score; S/R: Sarcomatoid/Rhabdoid; TCGA: The Cancer Genome Atlas;

**Figure S6:**
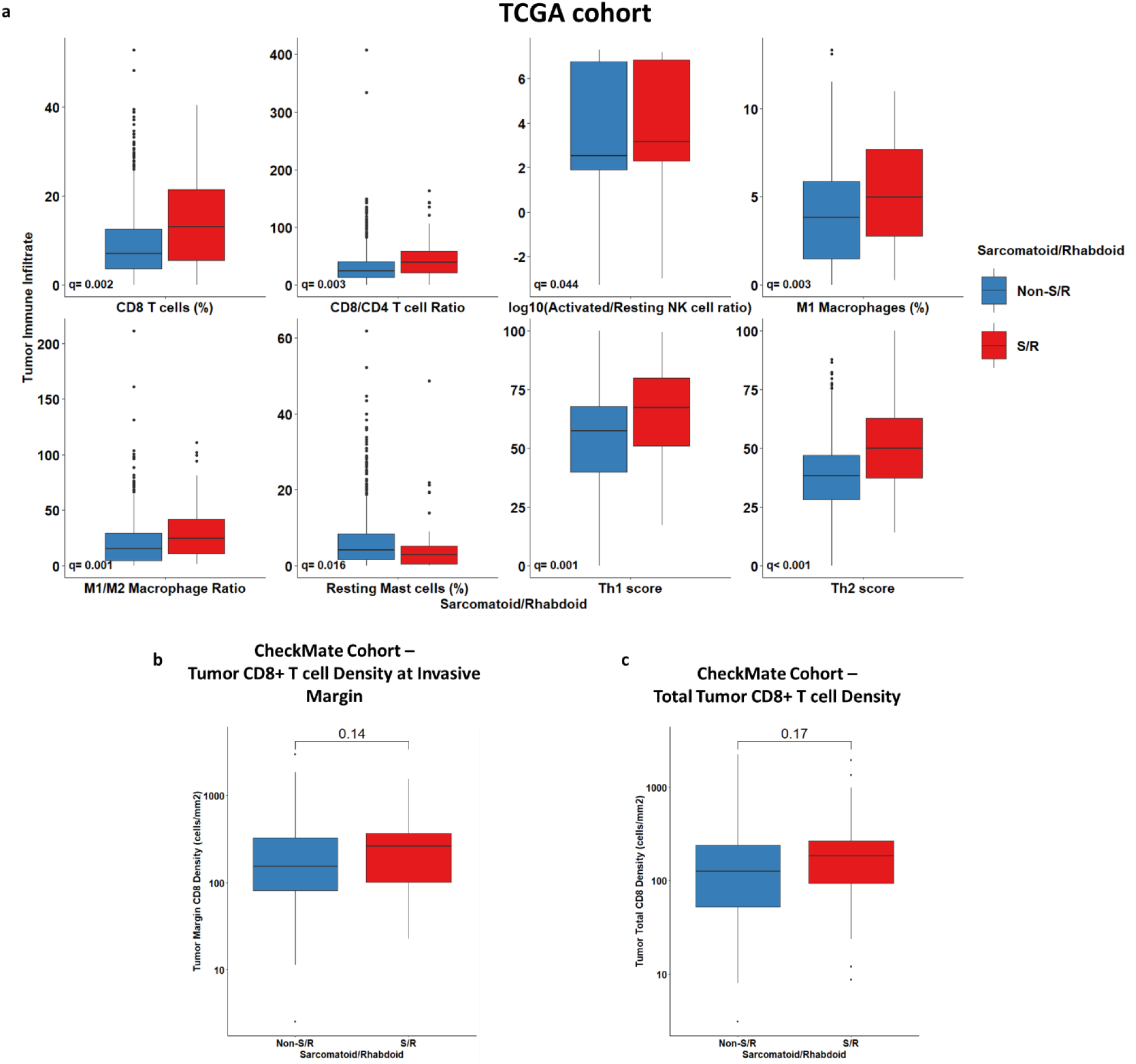
The improved outcomes of S/R RCC tumors on immune checkpoint inhibitors across clinical trial and real-word cohorts may be accounted for by an immune-inflamed phenotype (in relation to Fig. 4). (a) Boxplots of the comparison of CIBERSORTx and T helper immune cell populations between S/R and non-S/R RCC, with Mann-Whitney U test comparisons corrected for multiple comparison testing (q value reported). Only variables which were significant (q<0.05) in both the CheckMate and TCGA cohorts independently were shown. The TCGA results are displayed in this figure. Boxplots of the comparison of CD8+ T cell density at the (b) tumoral invasive margin and (c) throughout the tumor as determined by immunofluorescent staining in S/R compared to non-S/R RCC. Mann-Whitney U test p-values reported. S/R: Sarcomatoid/Rhabdoid; TCGA: The Cancer Genome Atlas.

**Figure S7:**
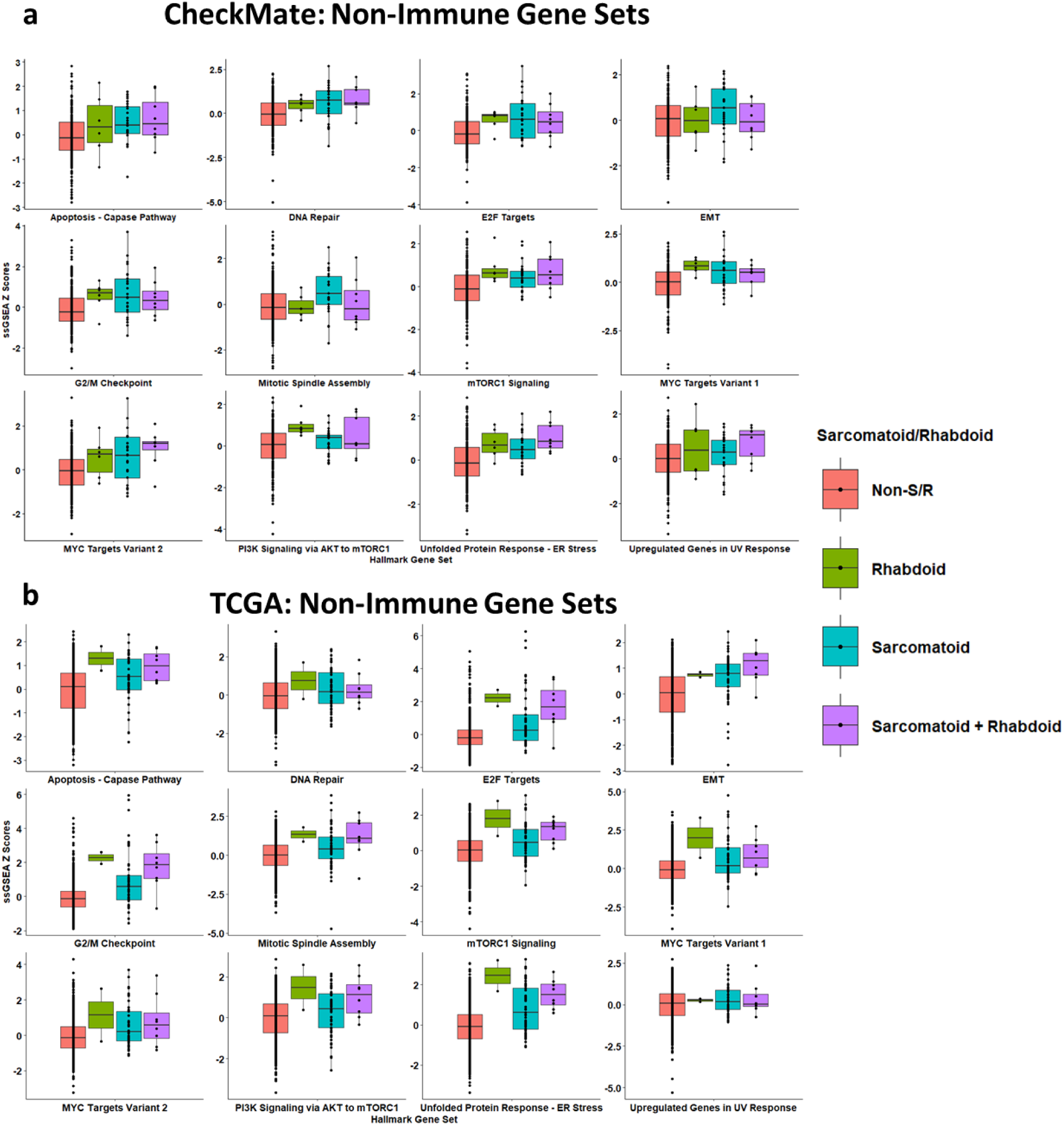
Breakdown of Z-score normalized ssGSEA scores in sarcomatoid, rhabdoid, and sarcomatoid and rhabdoid tumors of significantly enriched non-immune GSEA pathways in S/R RCC in the (a) CheckMate and (b) TCGA cohorts (in relation to Fig. 2). EMT: Epithelial Mesenchymal Transition; S/R: Sarcomatoid/Rhabdoid; ssGSEA: Single Sample Gene Set Enrichment Analysis

**Figure S8:**
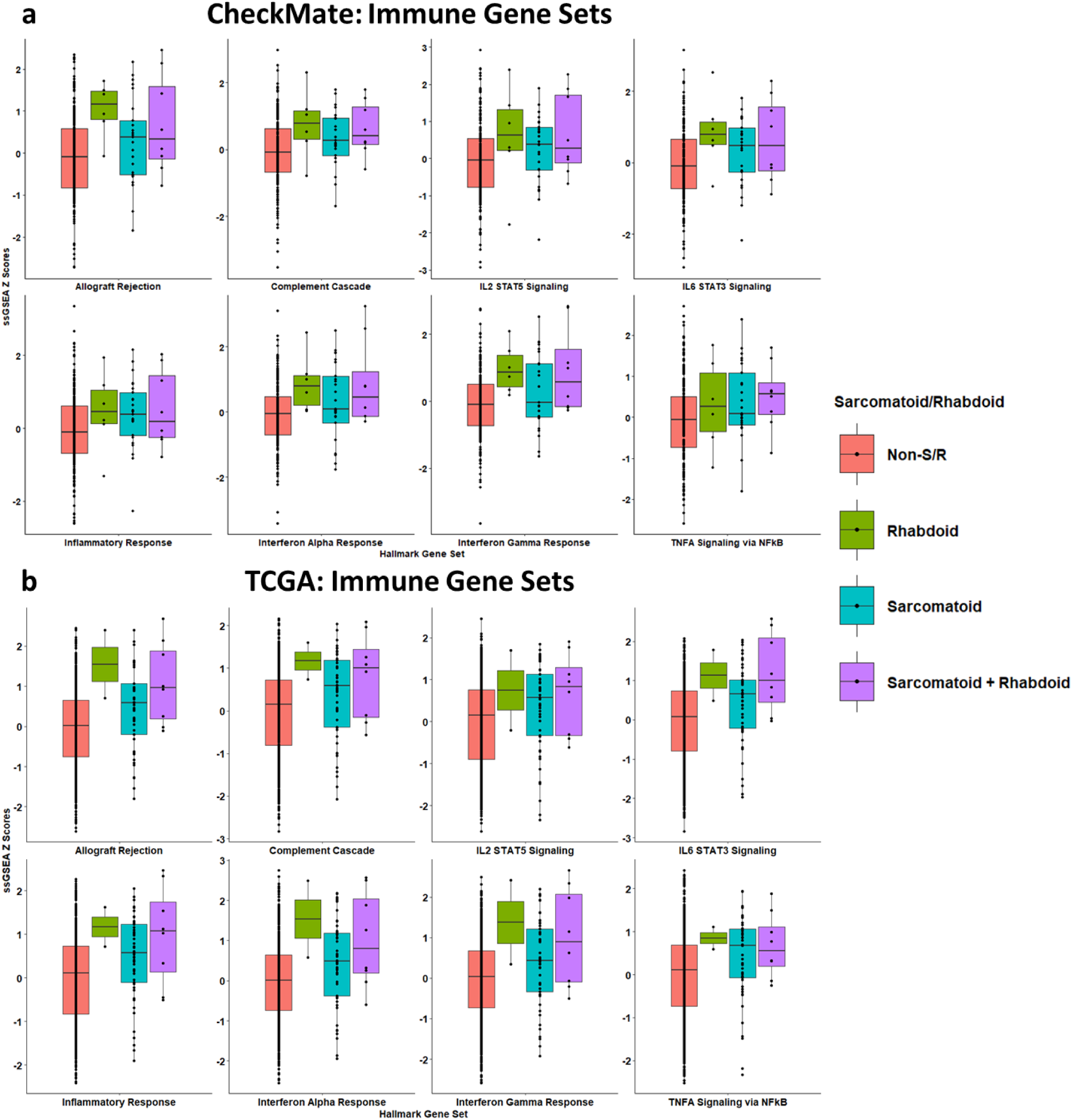
Breakdown of Z-score normalized ssGSEA scores in sarcomatoid, rhabdoid, and sarcomatoid and rhabdoid tumors of significantly enriched immune GSEA pathways in S/R RCC in the (a) CheckMate and (b) TCGA cohorts (in relation to Fig. 4). S/R: Sarcomatoid/Rhabdoid; ssGSEA: Single Sample Gene Set Enrichment Analysis

**Figure S9:**
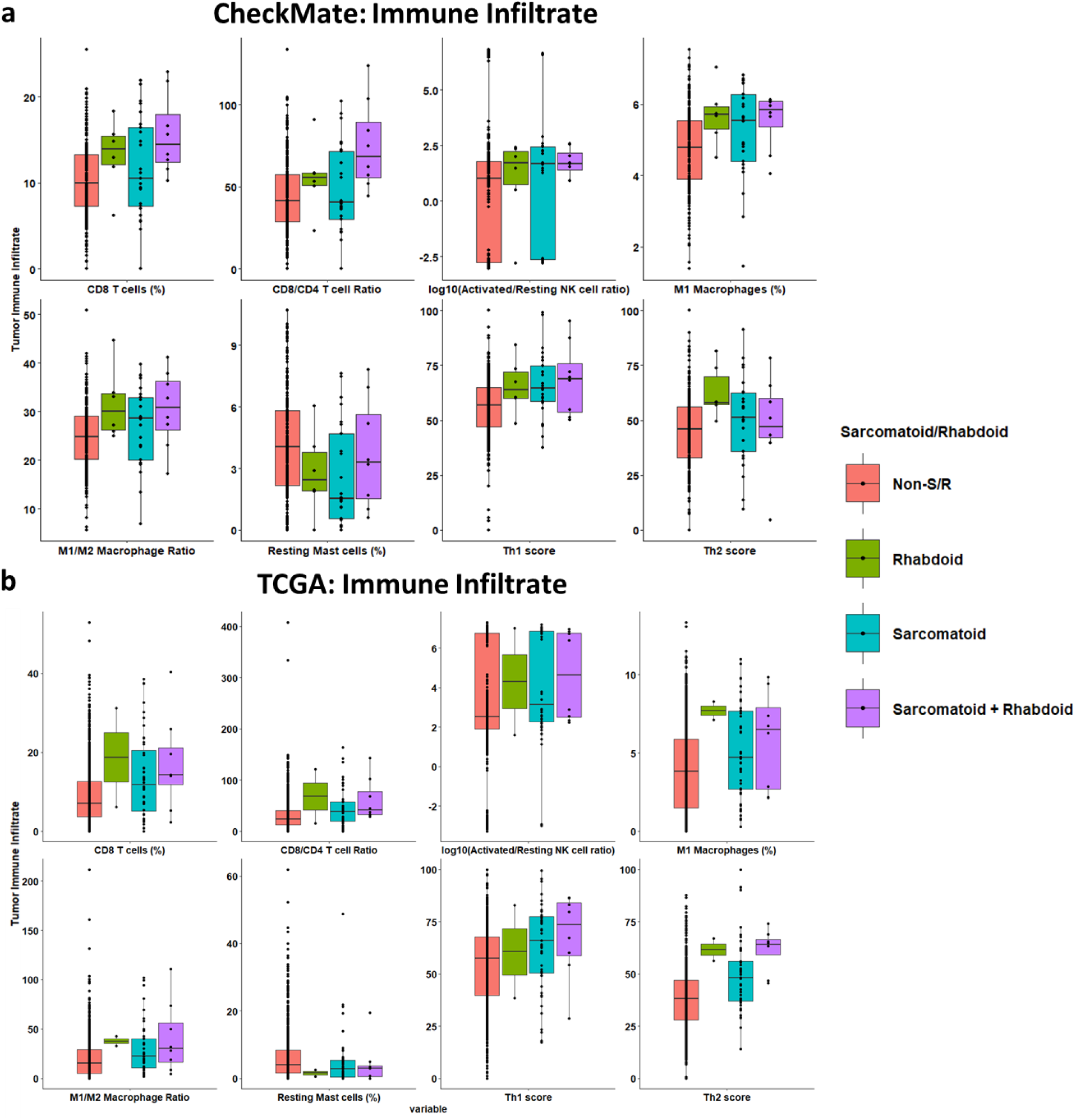
Breakdown of Z-score normalized ssGSEA scores in sarcomatoid, rhabdoid, and sarcomatoid and rhabdoid tumors of differentially enriched infiltrating immune cell populations in S/R RCC in the (a) CheckMate and (b) TCGA cohorts (in relation to Fig. 4). S/R: Sarcomatoid/Rhabdoid; ssGSEA: Single Sample Gene Set Enrichment Analysis

**Figure S10:**
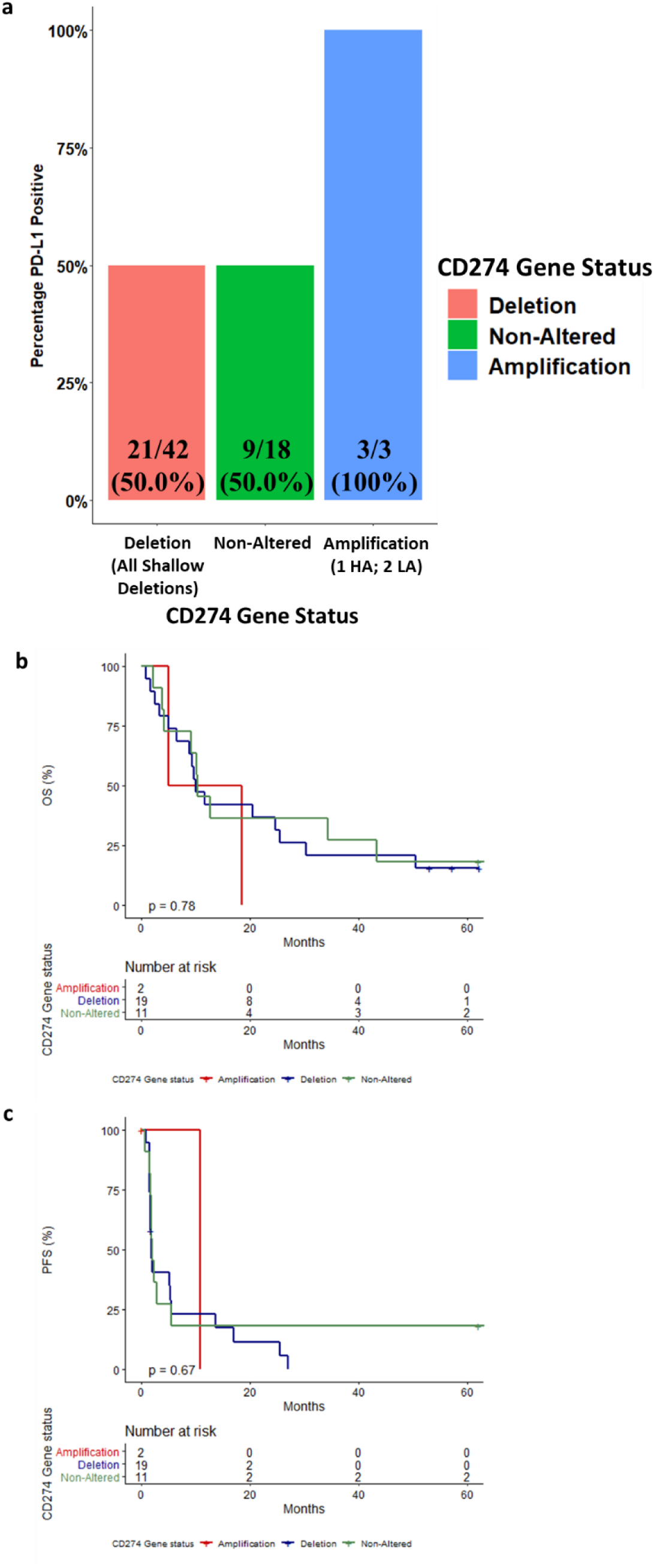
The improved outcomes of S/R RCC tumors on immune checkpoint inhibitors are not accounted for by *CD274* gene amplification. (a) Relationship between *CD274* (or PD-L1) gene status and PD-L1 expression in the subgroup of patients with S/R RCC that had WES and PD-L1 expression evaluated by IHC. Relationship between *CD274* (or PD-L1) gene status and survival outcomes on nivolumab in the subgroup of patients with S/R RCC that had WES and were treated by nivolumab; (b) OS and (c) PFS (in relation to Fig. 4). HA: High Amplification; LA: Low Amplification; OS: Overall Survival; PFS: Progression Free Survival.

**Figure S11:**
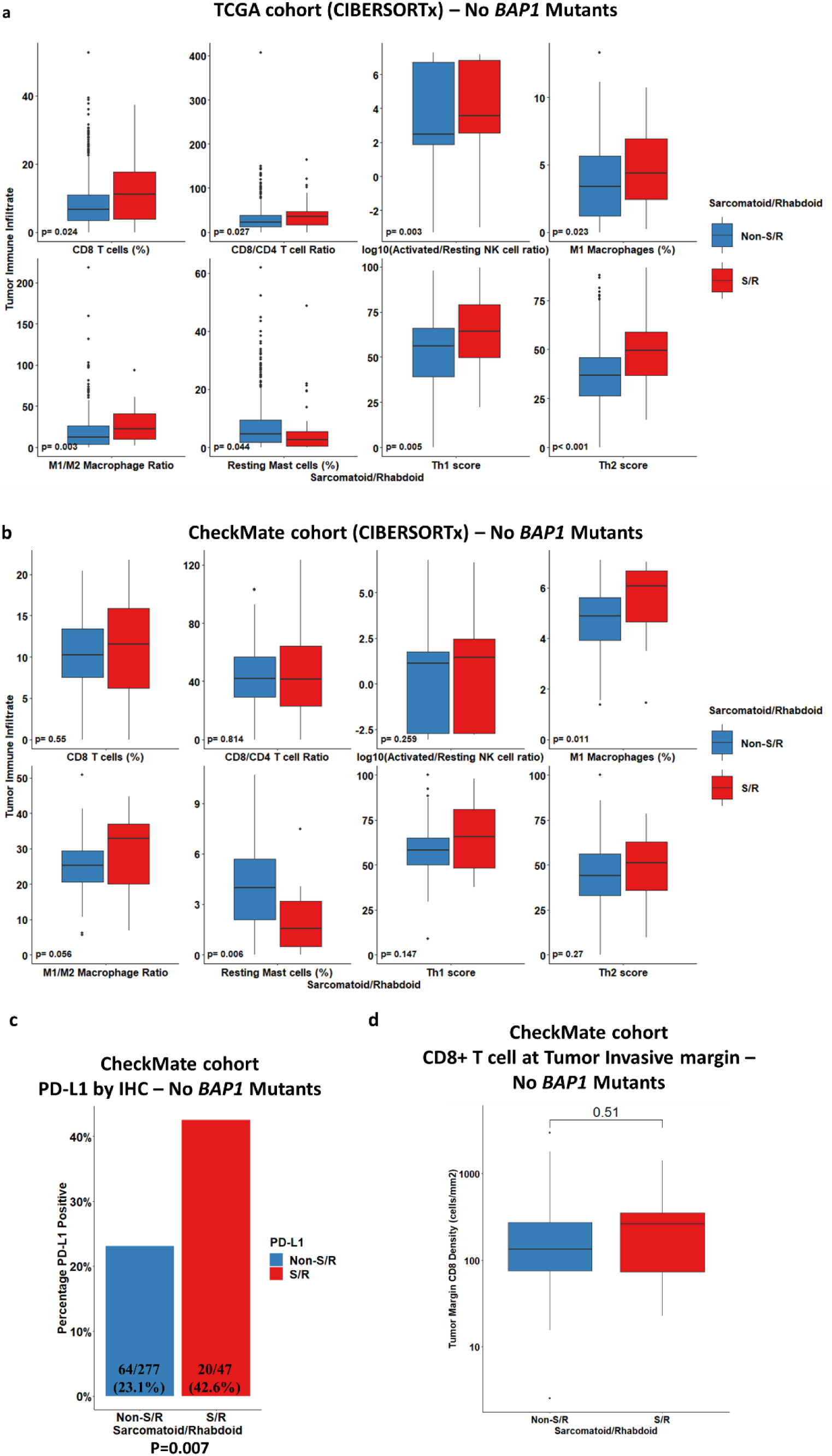
The immune-inflamed phenotype of S/R RCC tumors is independent of *BAP1* mutations. All plots exclude tumors with BAP1 mutations in both the S/R and non-S/R RCC groups (in relation to Fig. 4). Boxplots of the comparison of CIBERSORTx and T helper immune cell populations between S/R and non-S/R RCC, with Mann-Whitney U test (p-value reported) in the (a) TCGA and (b) CheckMate cohorts, excluding *BAP1* mutants. (c) Bar plot of the comparison of the proportions of tumors that were PD-L1 positive (≥1% on tumor cells) in S/R compared to non-S/R RCC, excluding *BAP1* mutants. Fisher’s exact test p-value reported. (d) Boxplot of the comparison of CD8+ T cell density at the tumoral invasive margin between S/R and non-S/R RCC, excluding *BAP1* mutants. Mann-Whitney U test p-value reported.

**Figure 12:**
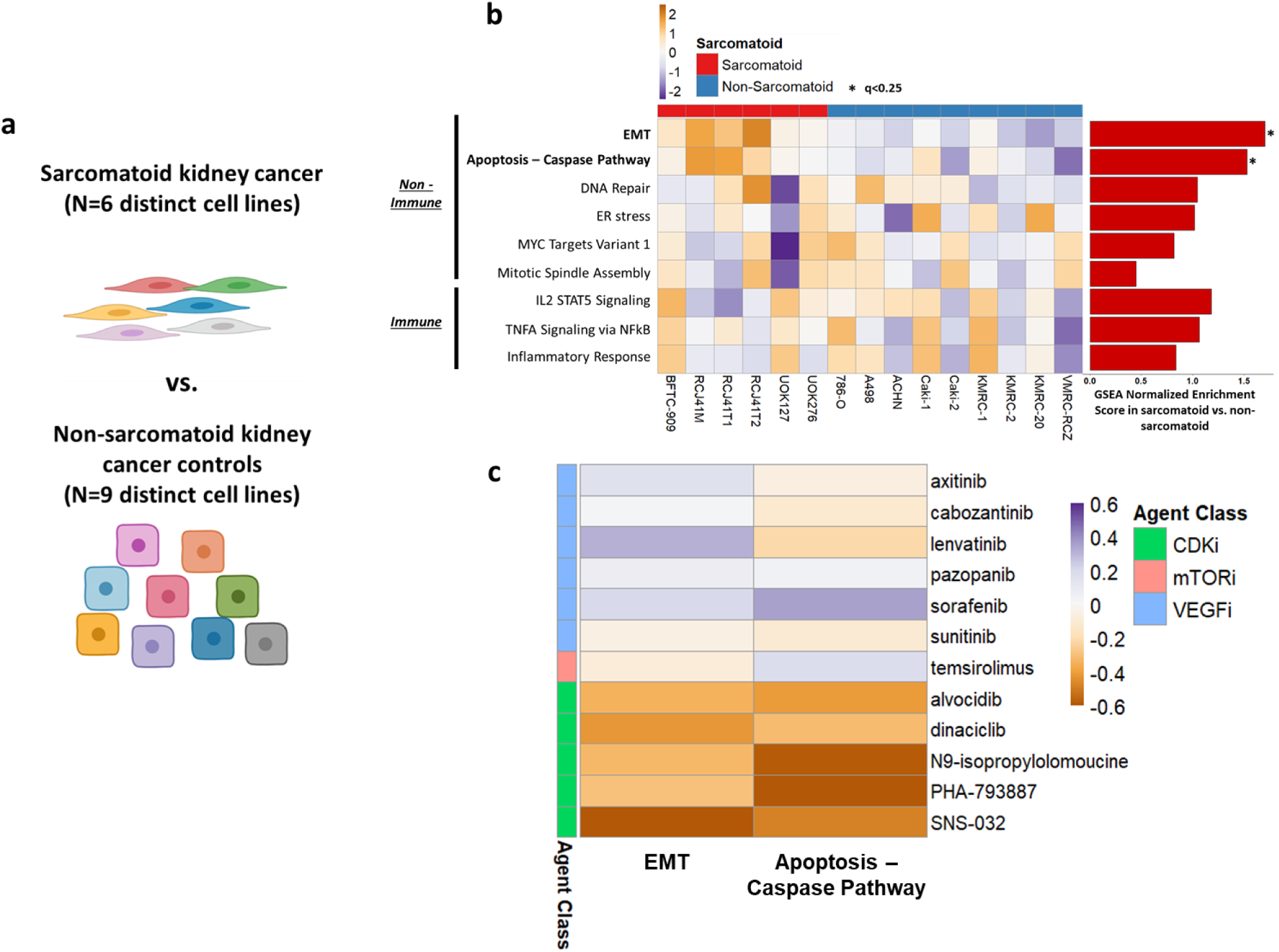
Baseline transcriptomic profiling of kidney cancer cell lines reveals that both immune and non-immune features of sarcomatoid tumors may be driven by the sarcomatoid component and suggests CDK as a potential therapeutic target. (a) GSEA was performed on the 50 “Hallmark” gene sets to compare 6 distinct sarcomatoid cell lines and 9 distinct non-sarcomatoid kidney cancer cell lines. (b) Heatmap and bar plot of the ssGSEA scores and GSEA normalized enrichment scores for the “Hallmark” gene sets that were found to be enriched in sarcomatoid compared to non-sarcomatoid cell lines. (c) Heatmap of the Pearson correlation coefficients between the area under curve (AUC) of the dose-response curve and the ssGSEA scores of the two pathways which were found to be significantly enriched in both cohorts of bulk RNA-seq and in the sarcomatoid cell lines (epithelial-mesenchymal transition and the apoptosis-caspase pathway). Agents are grouped by drug class and the color orange in this heatmap represents a negative correlation between ssGSEA score and AUC (indicating that a higher ssGSEA score correlates with greater drug sensitivity). The agents included in this figure are CDKi as well as the mTORi and VEGFi that are FDA-approved for metastatic renal cell carcinoma (for comparison). *q<0.25; CDKi: Cyclin-Dependent-Kinase Inhibitors; EMT: Epithelial Mesenchymal Transition; FDA: Food and Drug Administration; mTORi: Mammalian Target of Rapamycin Inhibitors; VEGFi: Vascular Endothelial Growth Factor Inhibitors.

**Figure S13:**
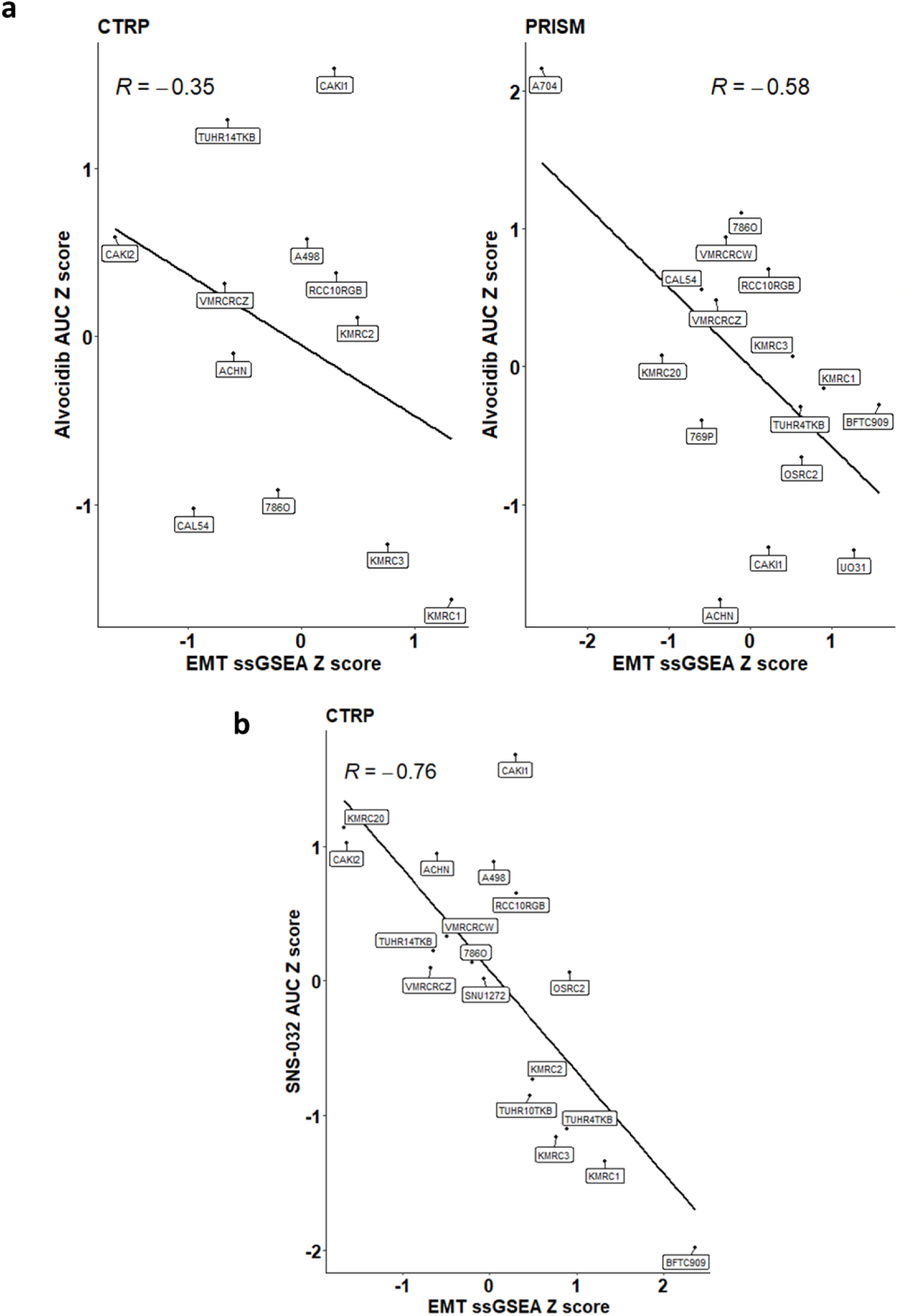
Scatter plots of correlations of transcriptomic characteristics of cell lines with areas under the curve of dose response curves in CTRP and PRISM for two CDK inhibitors (a) alvocidib and (b) SNS-032. Pearson r correlation coefficients shown. AUC: Area Under the Curve; EMT: Epithelial Mesenchymal Transition.

**Figure S14:**
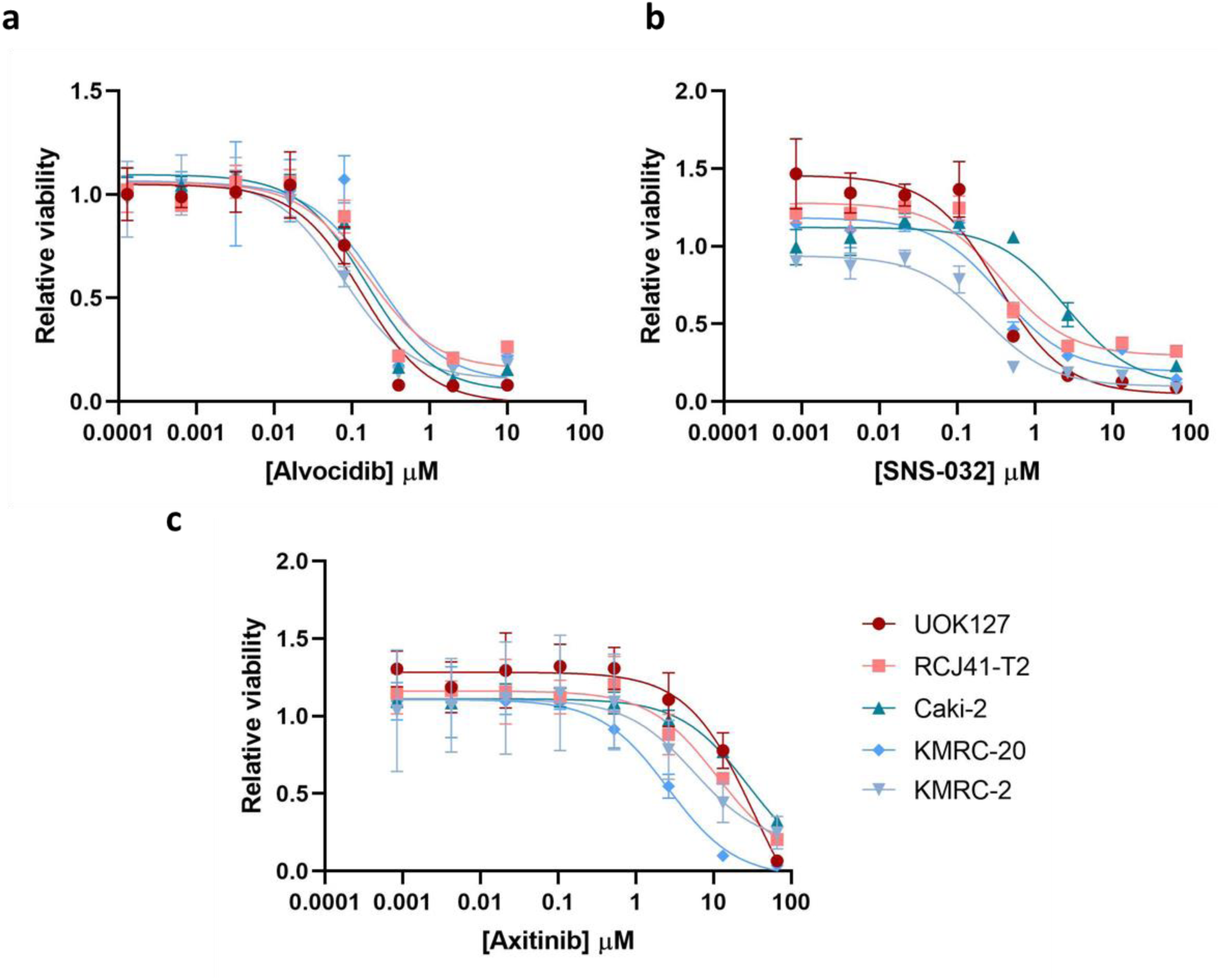
Dose-response curves of the in vitro cell line drug sensitivity assays for (a) alvocidib, (b) SNS-032, and (c) axitinib in two sarcomatoid cell lines (UOK 127 and RCJ41-T2) and three non-sarcomatoid cell lines (Caki-2, KMRC-20, KMRC-2).

**Figure S15:**
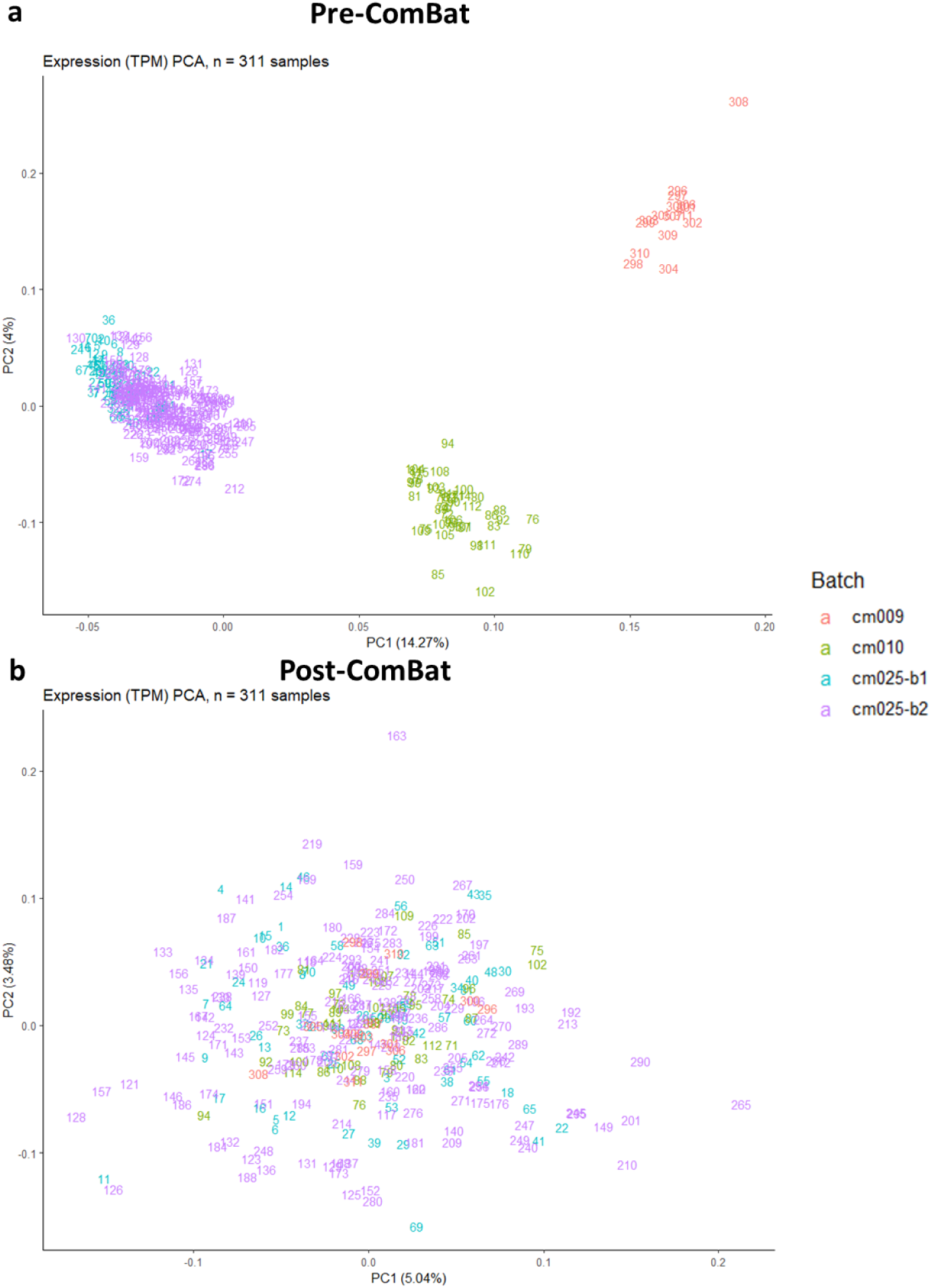
Principal component analysis plots of the UQ-normalized log2-transformed TPM matrix including the 3 known batches within the CheckMate cohort (a) pre-ComBat and (b) post-ComBat. cm010: CheckMate 010; cm-025-b1: CheckMate 025 Batch 1; cm-025-b2: CheckMate 025 Batch 2; PC1: Principal Component 1; PC2: Principal Component 2; PCA : Principal Component Analysis ; TPM : Transcripts-Per-Million.

## Supplementary Table Legends

Table S1: Baseline characteristics of the TCGA, CheckMate, and OncoPanel genomic cohorts, and clinical and genomic data of the OncoPanel cohort.

Table S2: Genomic analysis results of the TCGA, CheckMate, and OncoPanel genomic cohorts, genomic meta-analysis results, and breakdown of genomic alterations by background histology in the TCGA and OncoPanel cohorts.

Table S3: Gene level enrichment analyses of mutations in the OncoPanel cohort between epithelioid and S/R components of different S/R RCC tumors (Fisher’s exact tests) and in the Malouf cohort between epithelioid and S components of the same S RCC tumors (McNemar tests).

Table S4: Baseline characteristics of the TCGA and CheckMate RNA-sequencing cohorts.

Table S5: “Hallmark” and antigen presentation machinery gene set enrichment analysis results in the TCGA and CheckMate RNA-sequencing cohorts.

Table S6: “Hallmark” single sample gene set enrichment analysis in the TCGA and CheckMate RNA-sequencing cohorts and results of Cox regression analysis with overall survival.

Table S7: Gene-level differential gene expression analysis results (Mann-Whitney U test results) with log2 fold-changes of the mean. Genes that are significantly (q<0.05) upregulated or downregulated in the TCGA and CheckMate cohorts independently are also highlighted in separate tabs.

Table S8: Baseline characteristics of the Harvard, IMDC, and CheckMate clinical cohorts.

Table S9: CIBERSORTx deconvolution results in absolute mode of the CheckMate and TCGA cohorts with single sample gene set enrichment scores for Th1, Th2, and Th17 cells (scaled between 0 and 100) and Mann-Whitney U test comparison results in the TCGA and CheckMate cohorts independently.

Table S10: Baseline characteristics of patients that had their tumor tissue stained by immunohistochemistry for PD-L1 or CD8+ T cells by immunofluorescence.

Table S11: Raw and transformed TPM matrix of the 15 sequenced cell lines, quality control metrics by RNA-seqQC2, “Hallmark” gene set enrichment analysis of sarcomatoid vs. non-sarcomatoid cell lines, “Hallmark” single-sample gene set enrichment analysis of all 15 cell lines, epithelial-mesenchymal transition and apoptosis-caspase pathway single-sample gene set enrichment analysis of the 20 kidney cancer cell lines in CTRP v2 with drug sensitivity data, Pearson r correlation coefficients between single-sample gene set enrichment analysis scores and areas under the curve (AUC) of the dose-response curves for the 20 kidney cancer cell lines in CTRP v2 and in the PRISM secondary screen.

Table S12: Sarcomatoid and rhabdoid annotation for the TCGA KIPAN cohort.

Table S13: List of genes evaluated in the genomic analysis and table indicating which genes were included in each version of the OncoPanel assay

